# Whole-organism behavioral profiling reveals a role for dopamine in state-dependent motor program coupling in *C. elegans*

**DOI:** 10.1101/2020.03.27.011015

**Authors:** Nathan Cermak, Stephanie K. Yu, Rebekah Clark, Yung-Chi Huang, Steven W. Flavell

## Abstract

Animal behaviors are commonly organized into long-lasting states that coordinately impact the generation of diverse motor outputs such as feeding, locomotion, and grooming. However, the neural mechanisms that coordinate these diverse motor programs remain poorly understood. Here, we examine how the distinct motor programs of the nematode *C. elegans* are coupled together across behavioral states. We describe a new imaging platform that permits automated, simultaneous quantification of each of the main *C. elegans* motor programs over hours or days. Analysis of these whole-organism behavioral profiles shows that the motor programs coordinately change as animals switch behavioral states. Utilizing genetics, optogenetics, and calcium imaging, we identify a new role for dopamine in coupling locomotion and egg-laying together across states. These results provide new insights into how the diverse motor programs throughout an organism are coordinated and suggest that neuromodulators like dopamine can couple motor circuits together in a state-dependent manner.

## Introduction

As animals explore their environments, their nervous systems transition between a wide range of internal states that influence how sensory information is processed and how behaviors are generated (Anderson and Adolphs, 2014; Artiushin and Sehgal, 2017; Liu and Dan, 2019). These internal states of arousal, motivation, and mood typically alter many ongoing behaviors, impacting motor circuits that control diverse behavioral outputs such as feeding, grooming, and locomotion. A full understanding of how internal states are generated should explain how they coordinately alter the production of many complex motor outputs.

Neuromodulators play a central role in generating internal states. For example, neuropeptides like orexin and pigment dispersing factor (PDF) promote brain-wide states of wakefulness in mammals and flies, respectively (Saper et al., 2010; Taghert and Nitabach, 2012). In awake animals, norepinephrine controls arousal and attention (Aston-Jones and Cohen, 2005; Carter et al., 2010), the neuropeptides Tac2 and CRH induce states of heightened anxiety (Füzesi et al., 2016; Kormos and Gaszner, 2013; Zelikowsky et al., 2018), and the neuropeptides AgRP and NPY promote behaviors associated with hunger (Chen et al., 2019; Krashes et al., 2013). The anatomical organization of neuromodulatory systems makes them well-suited to impact a wide range of motor outputs: the projections of neuromodulator-producing neurons are typically very diffuse, targeting many brain regions. However, our mechanistic understanding of how neuromodulatory regulation of CNS circuits is propagated to diverse motor outputs remains limited.

In the simple nematode *C. elegans*, it should be feasible to determine how internal states influence every behavioral output of the animal. The *C. elegans* nervous system consists of 302 neurons, which act on 143 muscle cells (White et al., 1986). *C. elegans* generates a well-defined repertoire of motor programs, each with a devoted motor circuit and muscle group: locomotion, egg-laying, feeding, defecation, and postural changes of the head and body (de Bono and Maricq, 2005; Collins et al., 2016; Pirri et al., 2009; Schafer, 2005; Stephens et al., 2008). Quantitative studies of *C. elegans* locomotion have shown that animals switch between long-lasting behavioral states. In an extreme case, *C. elegans* animals cease all behaviors as they enter sleep-like states during development and after periods of stress (Nichols et al., 2017; Raizen et al., 2008; Van Buskirk and Sternberg, 2007). Awake animals in a food-rich environment switch between roaming and dwelling states, where they either rapidly explore the environment or restrict their movement to a small area (Ben Arous et al., 2009; Flavell et al., 2013; Fujiwara et al., 2002; Stern et al., 2017). After removal from food, *C. elegans* generates an area-restricted search state before switching to a dispersal state (Gray et al., 2005; Hills et al., 2004; López-Cruz et al., 2019; Wakabayashi et al., 2004). Like the internal states of mammals, these behavioral states in *C. elegans* last from minutes to hours and the transitions between states are abrupt. It remains unclear how the full repertoire of *C. elegans* behaviors is coordinated as awake *C. elegans* animals switch between behavioral states.

Neuromodulation in *C. elegans* plays a pivotal role in behavioral state control (Chase and Koelle, 2007; Li and Kim, 2008). Serotonin initiates and maintains dwelling states, while the neuropeptide PDF initiates and maintains roaming states (Choi et al., 2013; Flavell et al., 2013; Horvitz et al., 1982; Rhoades et al., 2019; Sawin et al., 2000; Stern et al., 2017). In addition, dopamine, tyramine, and octopamine influence behaviors associated with the presence or absence of food (Alkema et al., 2005; Chase et al., 2004; Horvitz et al., 1982; Sawin et al., 2000; Stern et al., 2017). The *C. elegans* sleep-like state is also strongly influenced by neuropeptides, including NLP-8, FLP-13, and FLP-24, and ligands for the NPR-1 receptor (Choi et al., 2013; Iannacone et al., 2017; Nath et al., 2016; Nelson et al., 2014). Command-like neurons that release neuromodulators are capable of driving state changes: the serotonergic NSM neurons induce dwelling states (Flavell et al., 2013; Rhoades et al., 2019) and the peptidergic ALA neuron evokes a sleep-like state (Hill et al., 2014; Nath et al., 2016; Nelson et al., 2014). However, there is not a one-to-one mapping between neuromodulators and specific states. For example, although serotonin release from NSM drives dwelling, serotonin release from ADF neurons has no apparent effect on dwelling and instead impacts other behaviors (Flavell et al., 2013). In addition, ALA releases at least three neuropeptides that have unique yet overlapping functions to inhibit downstream behaviors during sleep (Nath et al., 2016). However, each of these neuropeptides is also produced by other neurons that do not induce sleep. Thus, individual neuromodulators may exert different effects during different states and their combinatorial actions might give rise to the widespread behavioral changes that accompany each state.

In this study, we examine how neuromodulation coordinates diverse behavioral outputs to give rise to state-dependent changes in behavior. First, we describe a new imaging platform that permits simultaneous, automated quantification of each of the main *C. elegans* motor programs. We analyze hours-long behavioral recordings to fully characterize the behavioral states of this animal. We then identify a dopaminergic pathway that couples multiple motor programs together as animals switch behavioral states, showing that dopamine promotes egg-laying in a locomotion state-dependent manner. Our study provides new insights into how the diverse motor programs throughout an organism are coordinated and suggests that neuromodulators like dopamine can couple motor circuits together in a state-dependent manner.

## Results

### The distinct motor programs in *C. elegans* are coordinated with one another

To examine how the distinct motor programs generated by *C. elegans* animals are coordinated, we designed and constructed tracking microscopes and an accompanying software suite that permits simultaneous, automated measurement of numerous ongoing *C. elegans* motor programs (Figure 1A). The microscope, which shares several design features with previously described microscopy platforms (Yemini et al., 2013; see also Faumont and Lockery, 2006; Nguyen et al., 2016; Venkatachalam et al., 2016), collects brightfield images of individual animals (Figure 1B) at a frequency of 20 Hz and resolution of 1.4 um/pixel, which is sufficient to capture the most rapid and small-scale movements of *C. elegans*, such as pharyngeal motion (Avery and You, 2012; Lockery et al., 2012). A closed-loop tracking system reliably keeps animals in view over hours or days, while live data compression makes storage of approximately 1.7 million images per animal per day feasible. Although the parameters for data collection that we use here are tailored to *C. elegans* recordings, this low-cost, open-source microscopy platform should be useful for recordings of many small animals (see Methods for links to parts list and build tutorial).

**Figure 1.**
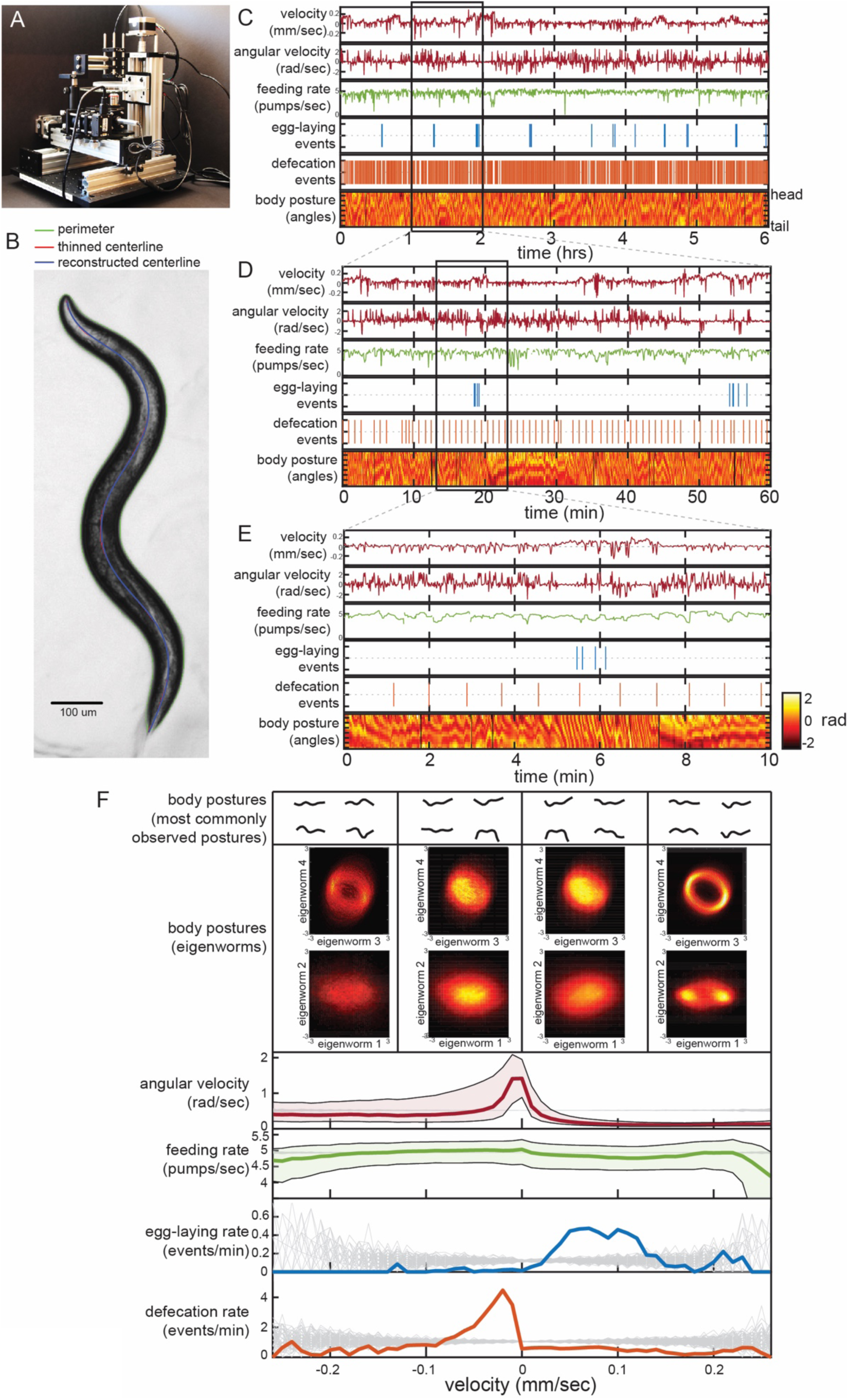
Simultaneous measurement of the diverse *C. elegans* motor programs. (A) Image of the tracking microscope. (B) Example image of a *C. elegans* animal from the tracking microscope. Green line denotes detected outline of worm; red line indicates worm’s centerline, obtained by thinning the thresholded image; blue line is the centerline as reconstructed from a spline-based 14-parameter representation. (C-E) Example dataset from the tracking microscope, showing the main *C. elegans* motor programs over 6 hours (C), 1 hour (D), or 10 min (E). (F) Average behaviors observed while animals travel at different velocities. Data are from 30 wild-type animals, where data points were binned based on instantaneous velocity. The average motor outputs in each bin are plotted. For angular velocity and feeding rate, data are shown as medians ± 25^th^ and 75^th^ percentiles. For egg-laying rate and defecation rate, data are shown as mean event rates (gray lines indicate random samples). For posture, data were segregated into four bins ranging from −0.25 mm/sec to 0.25 mm/sec. For each bin, we analyzed posture in two ways: by identifying and displaying the four most commonly observed postures (top), or by plotting 2D histograms of the four main eigenworms (middle). Eigenworms were calculated over the full datasets using previously described methods (Stephens et al., 2008).

We developed a software suite that allows us to automatically extract measurements of each of the main *C. elegans* motor programs from these video recordings. Because *C. elegans* is transparent, each movement of the animal – even movements inside the body, like those of the pharynx – is visible during brightfield imaging. Using this software suite, we are able to extract: body/head posture, locomotion, egg-laying, defecation, and pharyngeal pumping (i.e. feeding). Extracting body posture and locomotion are straightforward (Stephens et al., 2008) (Figure 1B; Figure 1—figure supplement 1A-D), but the detection of egg-laying, defecation, and pumping required us to implement tailored machine vision algorithms (Figure 1—figure supplement 1E-O; see Methods). To ensure the reliability of our measurements, we compared the automated scoring of egg-laying, defecation, and pumping to manually scored data and found a high level of concurrence (Figure 1—figure supplement 1G, J, O). These advances now allow us to extract near-comprehensive records of each animal’s behavior from hours-long recordings. The resulting datasets capture behavioral outputs over multiple timescales: from milliseconds-scale postural changes to hours-long behavioral states (Figure 1C-E shows an example dataset at multiple time resolutions).

To examine how the distinct motor programs of *C. elegans* are coordinated over time, we first performed simple analyses to relate the acute generation of each behavioral variable to the others. We examined data from 30 adult well-fed, wild-type animals, each recorded for six hours on a homogenous *E. coli* food source. To determine how each behavioral output varies as a function of locomotion, we examined average behavioral outputs across different animal velocities (Figure 1F). Egg-laying events were predominant during rapid forward locomotion, while defecation events were most commonly observed during paused or reverse movement. We examined typical body postures as a function of velocity by either plotting the most commonly observed postures at different velocities (Figure 1F, top) or by plotting 2D histograms of the eigenmodes of the angles along the body for data points when animals travel at different velocities (Figure 1F, middle) (Stephens et al., 2008). By both measures, we found that animal posture co-varies considerably with velocity, suggesting that animals sample different body postures at different velocities (Figure 1F). We also examined average behaviors that surround egg-laying and defecation events by plotting event-triggered averages (Figure 1—figure supplement 2). These analyses revealed many of the same relationships between the behavioral outputs, but also showed that egg-laying and defecation events each occur during stereotyped posture and locomotion changes, as has been previously reported (Collins et al., 2016; Hardaker et al., 2001; Nagy et al., 2015). We conclude from these analyses that there is extensive coordination between the distinct *C. elegans* motor programs.

### Characterization of behavioral states reveals a high incidence of egg-laying during roaming and multiple sub-modes of dwelling

To understand how these behaviors are coupled together over longer time scales, we sought to identify the long-lasting behavioral states that wild-type animals generate, so that we could determine which motor outputs are observed in each state. Unsupervised learning can reveal the underlying states that generate observed behavioral variables (Berman et al., 2014; Marques et al., 2018; Wiltschko et al., 2015). We and others have previously applied hidden Markov models (HMMs) to *C. elegans* locomotion parameters, which identifies roaming and dwelling states, as well as quiescence/sleep under certain conditions (Flavell et al., 2013; Gallagher et al., 2013). However, there may be a broader set of behavioral states that differ along other behavioral axes. To identify such states, we performed unsupervised discovery on the animal’s body posture (Figure 2A), which is the most complex behavioral variable and is coordinated with the other motor programs.

**Figure 2.**
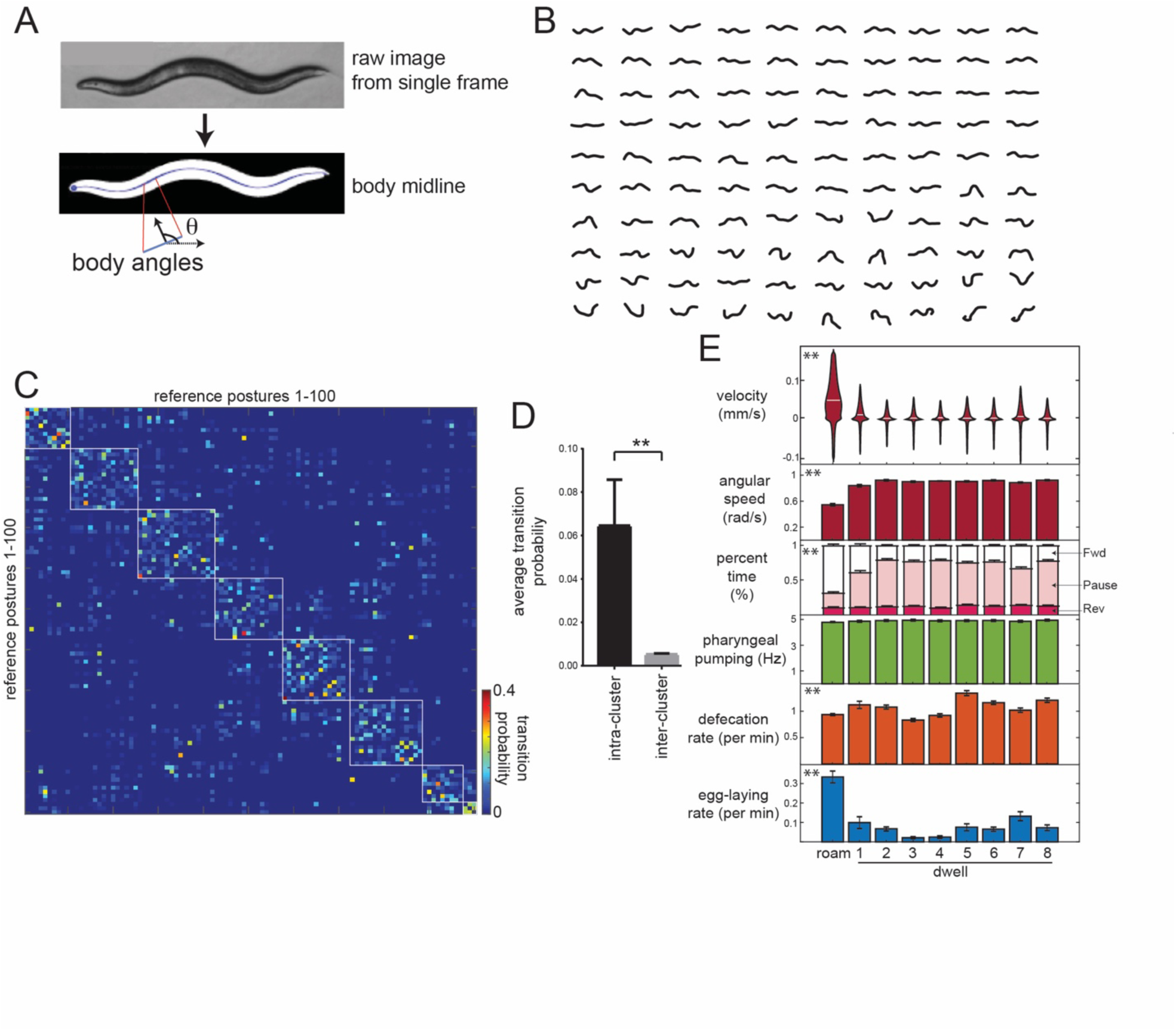
Identifying behavioral states through time series analysis of *C. elegans* posture. (A) Schematic showing that body posture is quantified as a vector of relative body angles, from head to tail. (B) Compendium of 100 reference postures that encompass the range of typical *C. elegans* postures. The compendium was derived by hierarchical clustering of many observed postures. (C) Transition matrix that shows probability of transitioning from each reference posture to the others. Self-transitions were excluded from this analysis. The rows of the matrix were clustered, and reference postures are sorted according to their cluster membership (eight clusters total). (D) Average transition rates between postures within the same cluster, versus average transition rates between postures in different clusters. **p<0.01, Wilcoxon test. (E) Average behaviors in each hidden state. Velocity is shown as a violin plot, while other behaviors are shown as averages across 30 wild-type animals. Data are means ± standard error of the mean (SEM). *p<0.0001, behaviors vary across states, ANOVA.

To describe postural changes in a compact manner, we used a previously described approach where we learned a compendium of reference postures that encompasses the broad range of body postures that animals display (Figure 2A-B) (Schwarz et al., 2015). To describe posture at any time point, we match the actual posture of an animal to its most similar match in the compendium. We then constructed a transition matrix that describes the probability of switching from each posture to the others (Figure 2C). Clustering the rows of this matrix revealed a striking, symmetric block-like structure, indicating that there are groups of postures, which we term “posture groups,” that animals transition back and forth between (Figure 2C-D; 11.9-fold higher transition rate to postures within the same group). We fit an HMM where the emissions are essentially these posture groups (see Methods). Based on Bayesian information criteria (BIC) estimates, a model with nine hidden states provided the best fit to the recorded data (Figure 2—figure supplement 1D; average state duration is 13 sec). Importantly, we reliably converged to the same model parameters, even when training was performed on different sets of animals and from different random starting conditions (Figure 2—figure supplement 1E-F). These results suggest that *C. elegans* alter their posture over slow time scales in a stereotyped manner.

We examined locomotion parameters in each of the nine hidden states. One of these behavioral states consisted of high forward velocity and low angular speed; based on these parameters, this state is equivalent to the previously defined roaming state (Figure 2E; labeled “Roam”) (Ben Arous et al., 2009; Flavell et al., 2013; Fujiwara et al., 2002). Animals traveled at lower velocities during the other eight states, but, in contrast to sleep states, they maintained high pumping and defecation rates in each (Figure 2E). To maintain consistency with previous literature, we describe these as dwelling sub-modes (Dwell1-8; see Video S1 for examples). Animals displayed significantly different postures in each of these sub-modes of dwelling (Figure 2—Figure Supplement 2). Whereas five of these states (Dwell2,3,4,5,8) reflected almost completely paused movement, Dwell6 involved more head and neck movements (Figure 2—Figure Supplement 2), Dwell1 involved more forward movement, and Dwell7 involved a higher rate of switching between forward and reverse locomotion (Figure 2E; Figure 2—figure supplement 1J-K). These data suggest that the previously defined dwelling state can be segmented into distinct sub-modes.

We also quantified the occurrence of other motor programs in each of these states (Figure 2E). Egg-laying rates were highest during roaming, whereas the dwelling sub-modes had smaller, but significant, differences in their egg-laying rates. This is consistent with previous observations that egg-laying is often accompanied by increased movement (Hardaker et al., 2001; McCloskey et al., 2017). Previous work has shown that animals display an acute reduction in speed immediately after egg-laying events (Hardaker et al., 2001). We found that this rapid speed change is present after egg-laying events during roaming and dwelling, but is far less pronounced when eggs are laid during roaming (Figure 2—figure supplement 1G-I). Defecation rates were also significantly different between the nine states, but feeding rates were the same across the states. Taken together, these data suggest that animals transition between a high-velocity/high-egg-laying roaming state and different sub-modes of dwelling in which they display distinct body postures and motor programs.

### Dopamine signaling promotes egg-laying during roaming states

We next sought to identify and characterize the neural circuits that allow animals to coordinate their motor programs across behavioral states. Here, we focused on the coupling between high-speed locomotion and egg-laying during the roaming state, since this was the most robust form of motor program coupling that we detected and our new microscopy platform allowed us to easily measure it. To test whether neuromodulation is critical for such coupling, we examined mutants lacking biogenic amine and neuropeptide neuromodulators.

For each mutant, we characterized the time spent in each behavioral state and the motor outputs within each state (Figure 3A). States were defined consistently across genotypes using parameters learned from wild-type animals. Serotonin-deficient *tph-1* mutants displayed reduced dwelling, high-speed roaming, reduced feeding, and fewer egg-laying events, as has been previously described (Figure 3A-B) (Avery and You, 2012; Flavell et al., 2013; Hobson et al., 2006; Horvitz et al., 1982). However, the increase in egg-laying rates during roaming states was largely intact, suggesting that serotonin is dispensable for this form of motor program coupling. *pdfr-1* mutants, deficient in PDF neuropeptide signaling, displayed a broad defect in roaming states: they spent less time roaming, traveled at lower velocity while roaming, and did not bias their egg-laying to the state as robustly as wild-type animals. Thus, although *pdfr-1* mutants are defective in locomotion and egg-laying during roaming, this may be part of a general deficit in their ability to display roaming states (Flavell et al., 2013), rather than a specific deficit in motor program coupling. *tdc-1* mutants that are defective in tyramine and octopamine displayed increased time in the roaming state. However, their egg-laying rates were still higher in the roaming state. *tbh-1* mutants defective in octopamine synthesis showed a mild increase in their roaming velocity, but no deficit in egg-laying. Overall, these mutants point to important roles for neuromodulation in regulating behavioral states in *C. elegans*, but do not provide insights into the coupling of egg-laying and locomotion during roaming.

**Fig. 3.**
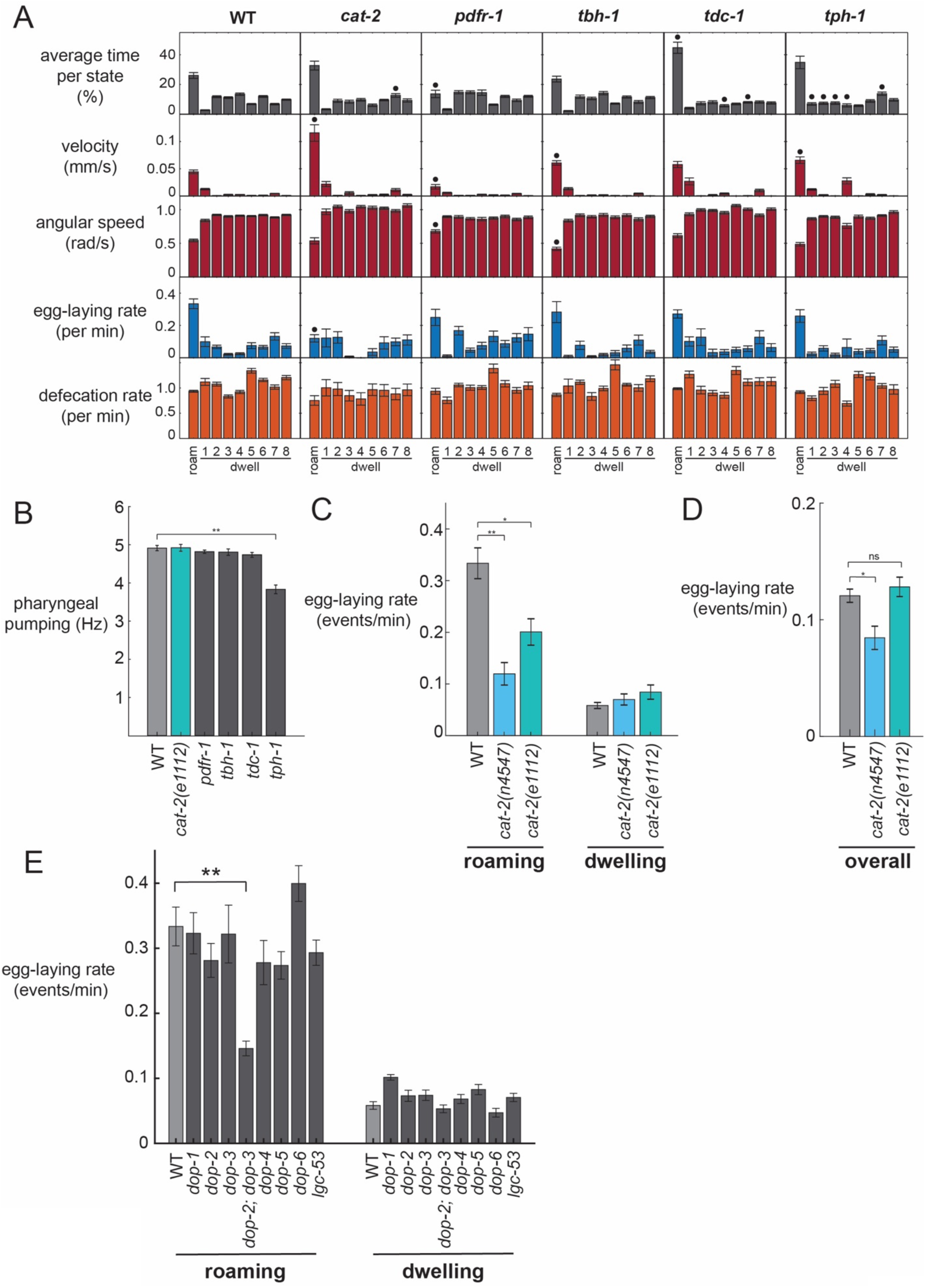
Analysis of neuromodulation mutants reveals a role for dopamine in state-dependent egg-laying. (A) Behavioral parameters of animals of the indicated genotypes across the posture-HMM states. Black dots indicate significant changes; *p<0.05, Bonferroni-corrected Welch’s t-test. The cat-2 mutant displayed here is *n4547* allele. (B) On food pharyngeal pumping rates in the neuromodulation mutants (data are pooled across all states). **p<0.001, Bonferroni-corrected Welch’s t-test. For (A-B), n=10-30 animals per genotype. (C) Egg-laying rates during roaming and dwelling states in two different dopamine-deficient *cat-2* null mutant strains. *p<0.01, Bonferroni-corrected Welch’s t-test. **p<0.001, Bonferroni-corrected Welch’s t-test. (D) Overall egg-laying rates in the *cat-2* null mutants. *p<0.05, Bonferroni-corrected Welch’s t-test. For (C-D), n=10-30 animals per genotype. (E) Egg-laying rates during roaming and dwelling for mutant animals lacking specific dopamine receptors. Data are means ± standard error of the mean (SEM). **p < 0.001, Bonferroni-corrected Welch’s t-test. n = 9-30 animals.

In contrast, dopamine-deficient *cat-2* mutants displayed a striking loss of coupling between locomotion and egg-laying. Although these animals displayed robust roaming states, their egg-laying rate was dramatically reduced during roaming (Figure 3A). In contrast, their egg-laying rate during the dwelling state was unaltered. *cat-2* animals travel at a higher velocity than wild-type animals while roaming (Figure 3A), but still display the same postures (Figure 3—figure supplement 1A), indicating that they are not broadly defective in the locomotor components of roaming (Sawin et al., 2000). Because they have unaltered egg-laying rates during dwelling, which is the more prevalent state, there is only a modest reduction in egg-laying overall in these mutant animals (Figure 3D). This suggests that they are not broadly defective in egg-laying, a finding that is consistent with the fact that they were not recovered as egg-laying defective (*egl*) in forward genetic screens. To ensure that this motor coupling phenotype was caused by the loss of the *cat-2* gene, we examined a second, independent null allele of *cat-2* and found that it caused a similar phenotype (Figure 3C–D). In addition, we rescued *cat-2* expression in the mutant via a transgene and found that this restored higher egg-laying rates during roaming (Figure 3—figure supplement 1B). These data suggest that dopamine is necessary for proper coupling between locomotion and egg-laying during roaming states and highlights the value of simultaneously quantifying multiple ongoing motor programs within the animal.

We considered whether the reduced egg-laying rates of *cat-2* mutants during roaming, but not dwelling, might be due to a floor effect, where the egg-laying rates during dwelling cannot be further reduced. To test for potential floor effects, we recorded mutant animals with deficits in the egg-laying motor circuit (*egl-1*). These animals showed a dramatic reduction in egg-laying during both roaming and dwelling states (Figure 3— figure supplement 1D), suggesting that floor effects do not account for the *cat-2* phenotype.

To further characterize the dopaminergic pathway that couples egg-laying and locomotion during roaming, we examined mutants lacking each of the seven known dopamine receptors in *C. elegans* (Figure 3E). None of these single mutants showed any egg-laying phenotypes. However, we found that mutant animals lacking the two D2-like dopamine receptors, *dop-2* and *dop-3*, displayed reduced egg-laying rates during roaming, but not dwelling, closely matching the *cat-2* mutant phenotype. These results suggest that *dop-2* and *dop-3* act together to regulate egg-laying during the roaming state.

In analyzing these mutant datasets, we considered whether the egg-laying phenotypes of the dopamine mutants could be due to an indirect effect, where increased forward velocity during roaming reduces egg-laying. However, an analysis of velocity and egg-laying rates across the receptors mutants suggests that this is not the case: *dop-2;dop-3* double mutants display normal velocities during roaming, but have dramatically reduced egg-laying rates during roaming (Figure 3—figure supplement 1C). This interpretation that the egg-laying effects are not due to locomotion changes is further corroborated by our optogenetics findings below. Altogether, these mutant analyses suggest that dopamine acts through the D2-like receptors DOP-2 and DOP-3 to promote egg-laying during the roaming state.

### Acute changes in dopaminergic neuron activity control state-dependent egg-laying

These genetic analyses are consistent with the possibility that dopamine signaling plays a direct role in elevating egg-laying rates during the roaming state. To examine whether dopamine acutely regulates egg-laying, we performed optogenetic studies to acutely alter the activity of the neurons that release dopamine. To acutely silence these neurons, we used the *dat-1* promoter (Flames and Hobert, 2009; Jayanthi et al., 1998) to drive expression of the GtACR2 anion channel (Govorunova et al., 2015) in all four dopaminergic cell types, CEPV, CEPD, ADE, and PDE (no single-neuron promoters have been described for any of these four cell types). Exposing these animals to light caused them to display lower egg-laying rates that persisted until the light was terminated (Figure 4A). We asked whether this reflected a reduction in egg-laying during roaming, dwelling, or both states by separately analyzing egg-laying rates during each of the states during the lights-on period (Figure 4B). This analysis indicated that the silencing of dopaminergic neurons reduced egg-laying rates while animals were roaming, but only had a mild, non-significant effect during dwelling. We examined the other behaviors automatically quantified by the tracking microscopes, but observed no significant effects of dopaminergic neuron silencing on feeding or defecation, and only a small, transient effect on locomotion (Figure 4—figure supplement 1A, 1F-H). These data corroborate our genetic findings and suggest that dopaminergic neurons function to acutely promote egg-laying in adult roaming animals.

**Fig. 4.**
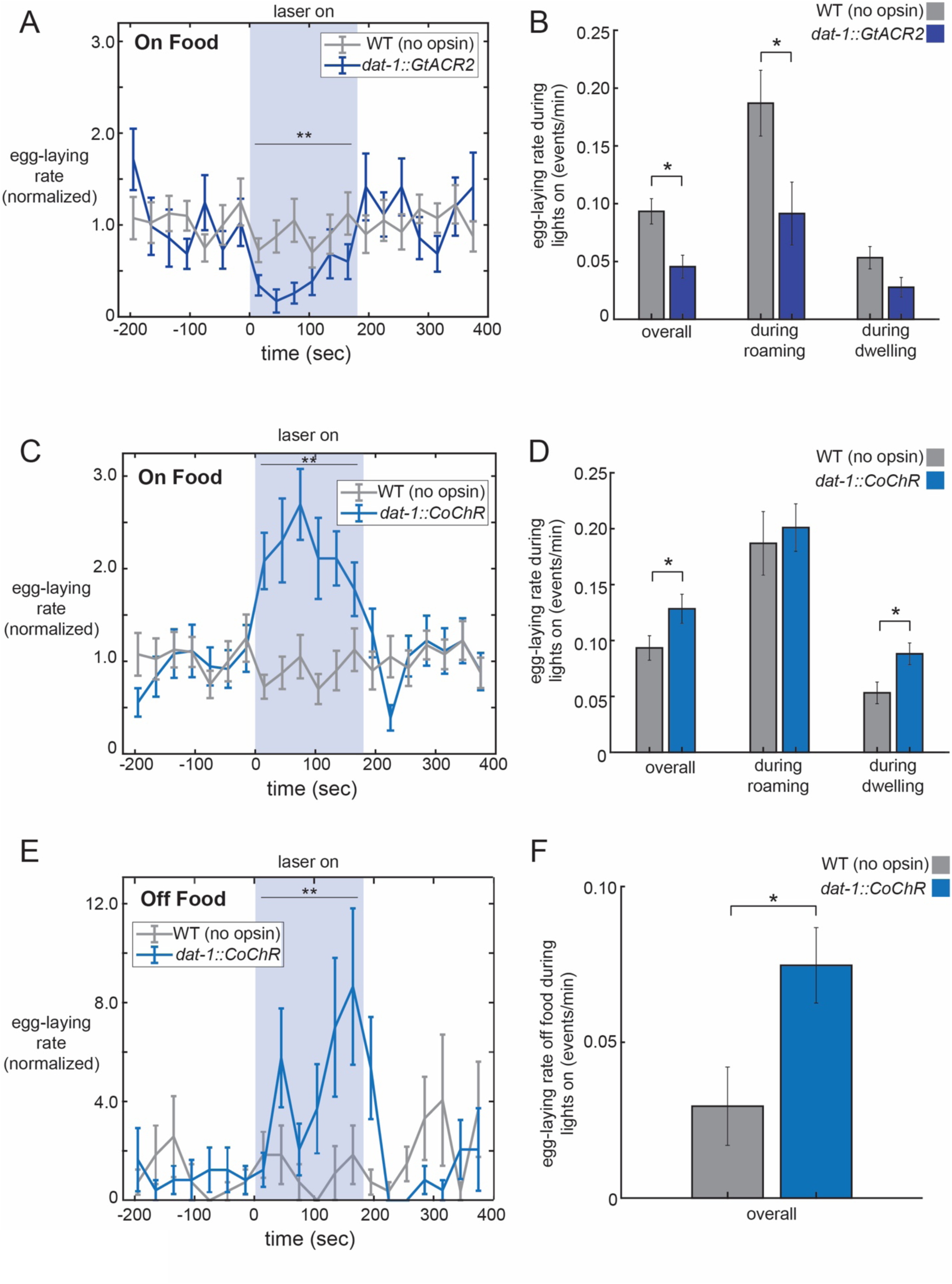
Optogenetic control of dopaminergic neurons alters egg-laying. (A) Inhibition of dopaminergic neurons via activation of *dat-1::GtACR2* reduces egg-laying rates. Data are shown as egg-laying rates, normalized to pre-stimulation baseline rates. Gray lines show wild-type animals stimulated with the same light pattern. **p<0.0001, paired t-test. (B) Average egg-laying rates during light exposure for *dat-1::GtACR2* and wild-type animals. Data are shown as overall rates, as well as rates during roaming and dwelling states specifically. *p<0.05, Welch’s t-test. For (A-B), n=416 light stimulation events across 12 animals for *dat-1::GtACR2*, and n=769 light events across 22 animals for wild-type. (C-D) Activation of dopaminergic neurons via activation of *dat-1::CoChR* increases egg-laying rates. Data are shown as in (A-B). **p<0.001, paired t-test. *p<0.05 Welch’s t-test. n=1221 light stimulation events across 35 animals for *dat-1::CoChR*, and n=769 light events across 22 animals for wild-type. (E-F) Effects of *dat-1::CoChR* activation in animals that are in the absence of food. Data are shown as in (A-B), except only overall egg-laying rates are shown since animals do not roam/dwell in the absence of food. *p<0.05, Welch’s t-test. **p<0.0001, paired t-test. n=387 light stimulation events across 44 animals for *dat-1::CoChR*, and n=327 light events across 38 animals for wild-type.

To examine whether exogenously increasing dopaminergic activity levels is sufficient to drive egg-laying, we acutely activated the four dopaminergic cell types. For these experiments, we used the *dat-1* promoter to drive expression of CoChR, a light-gated cation channel (Klapoetke et al., 2014). Activation of *dat-1::CoChR* for three minute durations caused animals to display elevated egg-laying rates (Figure 4C; minimal effects on velocity, Figure 4—figure supplement 1A, 1C). This increase in the rate of egg-laying persisted throughout the period of light exposure and was not necessarily time-locked to the moment of light onset. By contrast, acute activation of the egg-laying motor neuron HSN is known to trigger egg-laying events within seconds of light exposure (Emtage et al., 2012; Leifer et al., 2011). These data suggest that increased dopaminergic neuron activity increases the frequency of egg-laying events.

The observation that dopamine is necessary for proper egg-laying rates during roaming, but not dwelling, might be explained by elevated dopamine release during roaming. Alternatively, dopaminergic activity might be similar across states, but downstream circuits might integrate their detection of dopamine with a locomotion state variable, such that dopamine only enhances egg-laying during roaming. To begin to distinguish between these possibilities, we asked whether our externally-introduced, ectopic activation of dopaminergic neurons via optogenetics could increase egg-laying during roaming, dwelling, or both states. We found that light exposure to *dat-1::CoChR* animals elevated egg-laying during the dwelling state, but not during roaming (Figure 4D). The finding that the exogenous activation of dopaminergic neurons during dwelling is sufficient to increase egg-laying argues against a model where downstream circuits only respond to dopamine during roaming, but not dwelling. The finding that *dat-1::CoChR* activation does not further enhance egg-laying during roaming may reflect an occlusion effect, perhaps due to the already high egg-laying rates during roaming.

We again considered whether these effects of dopamine on egg-laying could be explained by an indirect effect on locomotion. Thus, we analyzed the velocity changes induced by optogenetic dopamine neuron silencing and activation. These two manipulations, which caused opposite effects on egg-laying, caused the same velocity change: a transient decrease in speed at light onset and a transient increase in speed at light offset (Figure 4—figure supplement 1A; includes no-opsin controls). Thus, the effects of these opposing optogenetic manipulations on egg-laying cannot be plausibly explained as an indirect consequence of altering locomotion.

Animals have dramatically lower egg-laying rates when they are removed from their food source. Because dopamine signaling is thought to be elevated in the presence of food (Sawin et al., 2000; Tanimoto et al., 2016), we asked whether increasing dopaminergic neuron activity in the absence of food was sufficient to increase egg-laying rates. Strikingly, we found that activation of *dat-1::CoChR* could drive animals to lay eggs in the absence of food at significantly higher rates (Figure 4E–F; velocity in Figure 4— figure supplement 1B). Altogether, these data suggest that native dopaminergic neuron activity in adult animals increases the probability of egg-laying during roaming states, and that exogenous dopaminergic neuron activity during dwelling or in the absence of food is sufficient to enhance egg-laying rates.

### Calcium dynamics in dopaminergic PDE neurons are phase-locked to egg-laying during roaming states

The above data suggest that dopamine release can enhance egg-laying and that native dopaminergic signaling in wild-type animals primarily promotes egg-laying during the roaming state. One possible explanation for these observations is that dopamine release might predominate during high-speed roaming. Thus, we next examined the native activity patterns of dopaminergic neurons in freely-moving animals. We constructed a transgenic strain expressing GCaMP6m under the *dat-1* promoter and recorded GCaMP signals in freely-moving animals using widefield imaging, as has been previously described (Flavell et al., 2013; Rhoades et al., 2019). The CEPD, CEPV, and ADE classes of neurons appeared to have static calcium levels while animals explored food and did not display dynamics that were dependent on locomotion speed (Figure 5— figure supplement 1A). However, the dopaminergic PDE neurons displayed robust calcium dynamics as animals freely explored a food lawn (Figure 5B). PDE neurons have short ciliated sensory dendrites that protrude through the cuticle along the dorsal side of animal in the posterior section of the body. The PDE axon travels along the ventral nerve cord, from the posterior to the anterior end of the animal. Notably, PDE is the only dopaminergic neuron whose neurite passes in close proximity to the egg-laying circuit and vulval muscles.

**Fig. 5.**
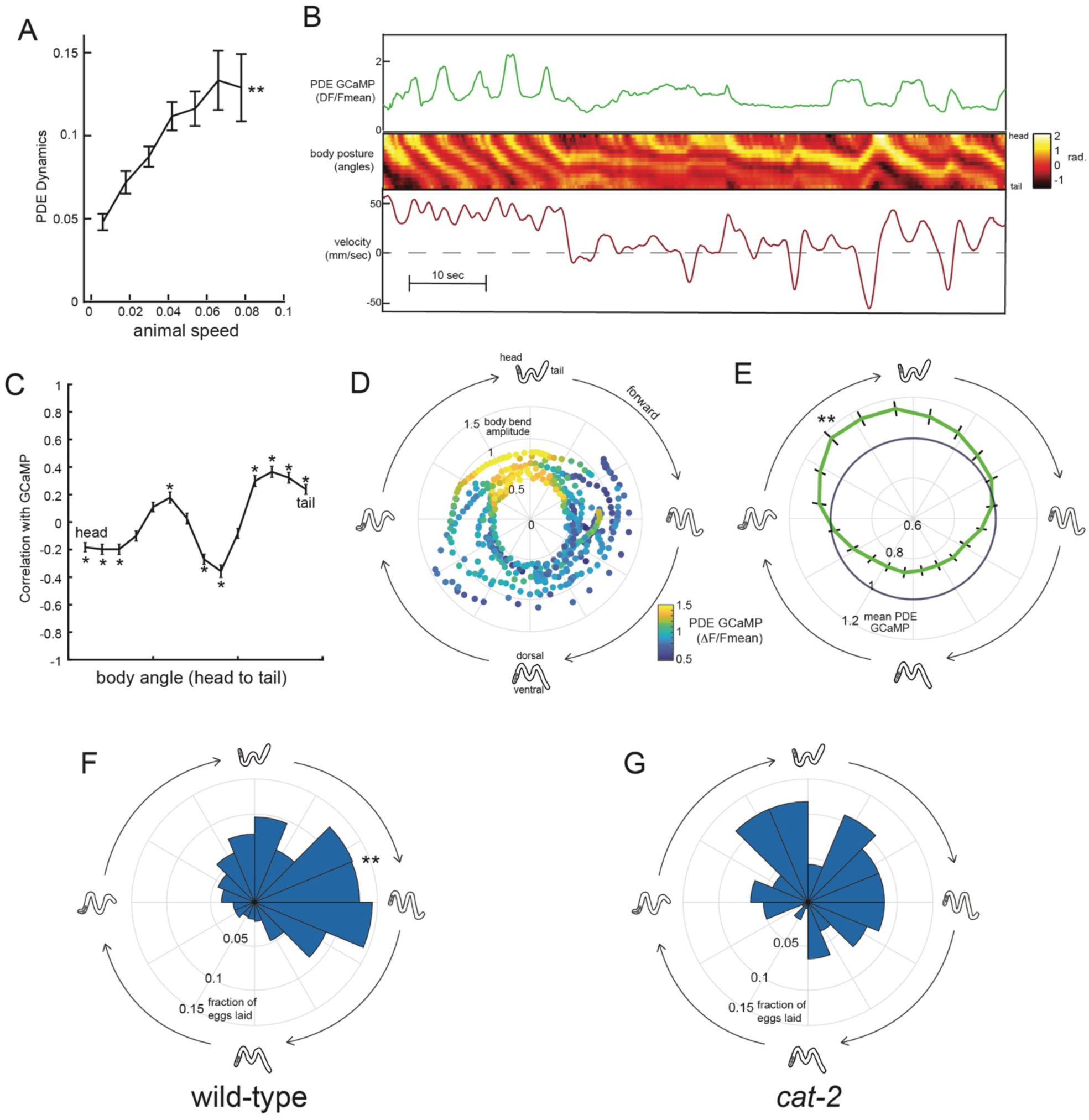
Dopaminergic PDE neurons display activity patterns phase-locked to egg-laying during roaming. (A) PDE dynamics increase with animal speed. PDE dynamics here is defined as the absolute value of the time derivative of the PDE GCaMP signal. **p<0.001, permutation test. (B) Example dataset from a wild-type animal, showing PDE::GCaMP6m signal, body posture (shown as body angles, from head to tail), and animal velocity. (C) Correlation coefficient of PDE activity with each of the 14 body angles is shown. *p<0.05, permutation test. (D) Example dataset showing how PDE activity (indicated by color) changes as animals proceed through stereotyped forward propagating bends during roaming. Theta values on the polar plot correspond to the phase of the forward propagating bend; radius corresponds to the depth of the body bends (which was quantified as the standard deviation of the mean-subtracted body angles). Corresponding body postures and the direction of the trajectories during forward movement are indicated. Note that PDE activity reliably increases during a specific phase of the forward propagating bend. (E) Average PDE activity at different phases of the forward propagating bend. Theta values are defined as in (D) and the radius indicates the mean PDE GCaMP signal. **p<0.001, paired-t-test. (F) Histogram depicting frequency of egg-laying events for wild-type animals at different phases during the forward-propagating bend. **p<0.001, Rayleigh z test. (G) Histogram depicting frequency of egg-laying events for *cat-2* mutants at different phases during the forward-propagating bend. *p<0.05 versus wild-type, Fisher’s exact test. For (A), (C), and (E), n=39 animals. For (F) and (G), n= 30 and 10 animals, respectively.

To quantify how PDE activity changes during locomotion, we first quantified the extent to which PDE dynamics depend on locomotion. We found that fluctuations in PDE calcium levels were significantly increased during high-speed roaming, which was not observed for the other dopaminergic cell types (Figure 5A; Figure 5—figure supplement 1A). As animals roamed, PDE activity oscillated and was highly correlated with the curvature of the animal’s body: PDE was most active when the dorsal body wall muscles were contracted on the posterior end of the body (Figure 5B shows an example; Fig. 5C shows correlations). When animals are moving forwards during roaming, repeating waves of muscle contractions pass through the body from head to tail. Thus, one result of the correlation between PDE activity and body curvature is that PDE is most active during a specific phase of each sinusoidal propagating wave (Figure 5D shows an example; Fig. 5E shows averages). Because PDE has an exposed sensory ending, we examined whether this activity pattern was impacted by the presence of bacterial food in the environment. Indeed, we found that PDE displayed significantly reduced dynamics during forward locomotion in the absence of food (Figure 5—figure supplement 1B). These relationships between PDE fluorescence and body posture were not observed when imaging PDE::GFP (instead of GCaMP), indicating that they were not due to motion artifacts (Figure 5 – figure supplement 1C-D). These data indicate that PDE calcium levels oscillate during forward movement on food, with a reliable activity peak during a specific phase of the forward propagating wave.

Previous studies suggested that egg-laying events also depend on body curvature (Collins et al., 2016), so we also quantified the frequency of egg-laying events during each sinusoidal propagating wave of muscle contractions during roaming. Indeed, egg-laying events were strongly biased to a specific phase of the sinusoidal wave (Figure 5F). Notably, the phase of maximal PDE activity overlapped with and shortly preceded the phase of maximal egg-laying. This relationship raised the possibility that dopamine release during roaming might not only promote egg-laying events overall, but might also bias them to occur during a specific phase of the sinusoidal wave. To test this possibility, we examined the distribution of egg-laying events in dopamine-deficient *cat-2* mutants and found that there was a significant decrease in the phase-dependence of egg-laying (Figure 5G). This difference was not due to a general change in body postures during roaming, since wild-type animals and *cat-2* mutants displayed similar postures during forward propagating waves while roaming (Figure 5—Figure Supplement 1G-H). Together, these data suggest that the activity of dopaminergic PDE neurons is phase-locked to egg-laying events during roaming and that dopamine signaling is required for proper coordination of body posture and egg-laying during roaming.

### GABAergic signaling is required for dopamine-induced egg-laying

We next sought to identify the downstream circuits through which dopaminergic neurons act to elevate egg-laying rates. To identify these downstream components, we examined whether specific neurons or neurotransmitters were necessary for dopamine-dependent egg-laying. Egg-laying in *C. elegans* requires contraction of the vulval muscles, which receive many synapses from HSN and VC neurons and a smaller number of synapses from cholinergic VA/VB neurons and GABAergic VD neurons. We optogenetically activated dopaminergic neurons in mutant animals lacking each of these four inputs: (1) *egl-1* mutants lacking HSNs, (2) *lin-39* mutants lacking VCs, (3) *acr-2* mutants with reduced cholinergic neuron activity, and (4) *unc-25* mutants with abolished GABAergic transmission (Figure 6A shows fold-change in egg-laying during lights-on, compared to lights-off). *dat-1::CoChR* activation still elevated egg-laying rates in *acr-2* mutants, suggesting that cholinergic transmission in ventral cord neurons is not essential for these effects. However, *dat-1::CoChR* activation had a reduced effect in *egl-1* and *lin-39* mutants and failed to elevate egg-laying rates in *unc-25* mutants. The finding that HSN and VCs are required for dopamine-induced egg-laying is not surprising, since these motor neurons are centrally involved in driving egg-laying, but the role of GABAergic signaling in egg-laying has not been well-studied, so we examined this interaction more closely.

**Fig. 6.**
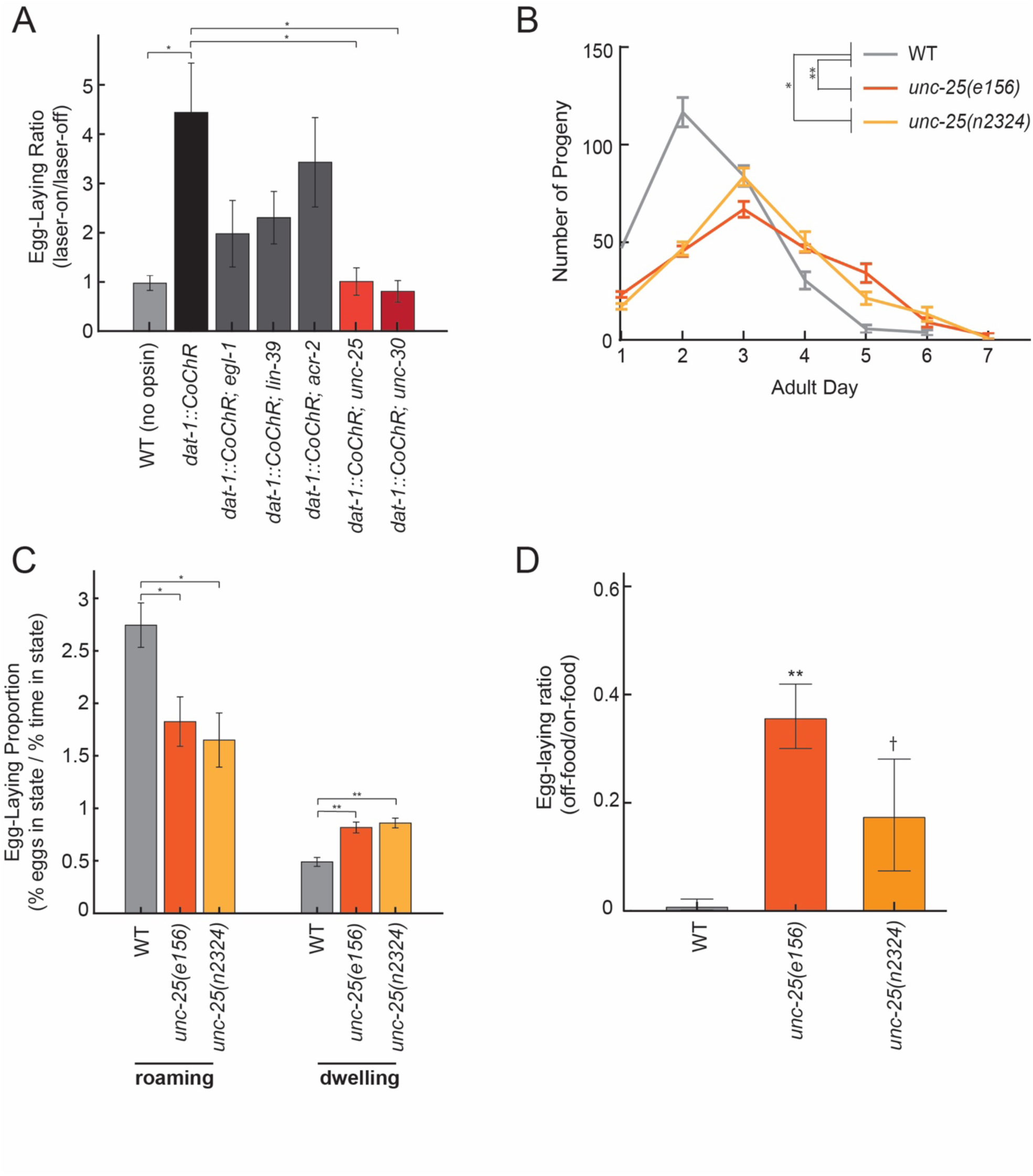
Dopamine elevates egg-laying in a GABA-dependent manner. (A) Effects of *dat-1::CoChR* activation in genetic mutants with disrupted components of egg-laying circuitry. Data are shown as the fold increase in egg-laying during lights-on period, compared to lights-off period (a value of 1 indicates no effect of the light on egg-laying). *p<0.01, Welch’s t-test. n= 9–35 animals per genotype. (B) Two independent null mutants lacking *unc-25*, which is required for GABA synthesis, have an altered profile of egg-laying. Data are shown as average eggs laid per day, with the first day being the first 24 hours after L4. *p<0.05 and **p<0.01, effect of genotype in two-factor ANOVA and post-hoc pairwise Dunnett test. n=5-10 animals per genotype. (C) Percent of eggs laid during roaming and dwelling for two independent null mutants of *unc-25*, which are defective in GABA synthesis. Here, we display data in this format to normalize for the reduced brood size in *unc-25*. *p<0.01, Welch’s t-test. **p<0.0001, Welch’s t-test. n= 11–30 animals per genotype. (D) Egg-laying in the absence of food is elevated in *unc-25* mutants. Data are shown as a ratio of eggs laid off food divided by eggs laid on food. **p<0.01 and †p<0.1 in ANOVA and post-hoc pairwise Dunnett test. n=5-6 plates per genotype, each with 10 animals per plate.

In *C. elegans*, GABA is produced by a small set of neurons in the head and tail, and by the VD/DD motor neurons in the ventral cord. To clarify which neurons mediate dopamine-dependent egg-laying, we activated *dat-1::CoChR* in *unc-30* mutants, which lack GABA in the VD/DD neurons, but not in the other GABAergic cell types in the head and tail (Figure 6A). We found that *dat-1::CoChR* failed to elevate egg-laying in *unc-30* mutants, indicating that GABA production by VD/DD neurons is critical for this effect.

A previous study showed that application of muscimol, a GABA_A_ receptor agonist, strongly inhibits egg-laying (Tanis et al., 2009), but the effects of reducing GABAergic signaling on egg-laying had not been previously examined. Thus, we performed experiments to clarify the role of native GABAergic signaling in egg-laying. Here, we compared wild-type animals to two independent *unc-25* null mutant strains, which both have attenuated GABA synthesis. Compared to wild-type animals, *unc-25* mutants had a slightly lower brood size (∼220 eggs per animal vs. ∼280 in wild-type) and laid their eggs over their first 5-6 days of adulthood, versus 3-4 days in wild-type animals (Figure 6B). As a result, during their first day of adulthood *unc-25* mutants displayed lower overall egg production, but then they displayed increased egg production as five- and six-day old adults (Figure 6B). We examined state-dependent egg-laying in these animals and found that, after adjusting for their lower egg production, one-day old *unc-25* mutants displayed a relative increase in egg-laying during dwelling, as compared to roaming (Figure 6C). To test whether GABA also regulates egg-laying across environmental conditions, we examined egg-laying in the absence of food and found that *unc-25* mutants displayed an elevated egg-laying rate off food, compared to wild-type animals (Figure 6D). Altogether, these results indicate that native GABAergic signaling regulates egg-laying, specifically playing an inhibitory role during dwelling and in the absence of food. This inhibitory role is consistent with the previous finding that muscimol inhibits egg-laying (Tanis et al., 2009). Given that GABAergic signaling is required for dopamine-dependent egg-laying, one possible explanation for these effects is that dopamine release may inhibit GABAergic neurons that function to inhibit egg-laying.

## Discussion

Behavioral states impact the generation of diverse motor outputs, but the neural mechanisms that allow them to exert these widespread effects are poorly understood. To examine this problem at whole-organism scale, we developed a new method that permits simultaneous, automated quantification of diverse *C. elegans* motor programs over long time scales. Analysis of these near-comprehensive records of animal behavior show that there is extensive coordination between the different motor programs of this animal. By combining this approach with genetics and optogenetics, we uncovered a new role for dopamine in promoting egg-laying in a locomotion state-dependent manner. Our results provide new insights into how the diverse motor programs throughout an organism are coordinated, and suggest that neuromodulators like dopamine can couple motor circuits together in a state-dependent manner.

To understand how internal states are represented in the brain, it is critical to obtain a quantitative description of the full repertoire of behaviors that are influenced by a given state. The transparency of *C. elegans* allows us to observe each of the motor actions of this animal, even the movements of internal muscle groups like the pharynx. Thus, by using tracking microscopes and machine vision software, we simultaneously quantified the diverse motor programs of this animal. These datasets revealed widespread coordination among distinct motor programs. For example, roaming and dwelling states that were previously described based on locomotion parameters also show robust differences in other motor programs, like egg-laying. Moreover, dwelling states can be segmented into different sub-modes based on reliable differences in body posture and other motor programs. We note that so far our analyses have been limited to well-fed adult animals exploring a homogeneous *E. coli* food source. The structure of *C. elegans* behavior could be quite different when animals are different ages (Stern et al., 2017), exposed to different stimuli, or in different physiological states. Several factors likely contribute to the coupling between diverse motor programs: changing internal state variables and recruitment of neuromodulators (Donnelly et al., 2013; Flavell et al., 2013; Pirri et al., 2009; Raizen et al., 2008; Van Buskirk and Sternberg, 2007), corollary discharges and related motor signals that allow motor circuits to interact (Gordus et al., 2015; Liu et al., 2018), and proprioceptive or environmental feedback (Goodman and Sengupta, 2019; Hu et al., 2011; Li et al., 2006; Wen et al., 2012; Yeon et al., 2018). Combining the whole-organism behavioral profiling approach described here with recently-developed whole-brain calcium imaging approaches (Kato et al., 2015; Nguyen et al., 2016; Venkatachalam et al., 2016; Yemini and Hobert, 2019) could yield a more complete understanding of which neural mechanisms are at work.

To begin to clarify neural mechanisms that underlie coordination between motor programs, we investigated a particularly robust form of motor program coupling: the increased frequency of egg-laying during high-speed roaming states. In wild-type animals exposed to a food source, egg-laying rates are approximately six-fold higher during roaming as compared to dwelling. This strategy might allow animals to distribute their eggs throughout a food resource. We found that dopamine signaling was required for this coordination: mutants with reduced dopamine levels had lower egg-laying rates during roaming states, but not during dwelling. Previous work has shown that native dopamine signaling in *C. elegans* is involved in driving slow locomotion in response to a bacterial food lawn, an effect called the basal slowing response (Sawin et al., 2000). Indeed, we observed increased forward velocity in *cat-2* mutants (Figure 3A). However, multiple lines of evidence suggest that the effects of dopamine on egg-laying appear are separable from the locomotion effects: (1) *dop-2; dop-3* dopamine receptor mutants have reduced egg-laying during roaming, but normal roaming velocities (Figure 3E; Figure 3—figure supplement 1C), and (2) optogenetic silencing and activation of dopaminergic neurons have opposite effects on egg-laying, even though they have the same modest effect on locomotion. We have not yet mapped out where the *dop-2* and *dop-3* receptors function to control egg-laying, though it is intriguing that *dop-3* is known to be expressed in VD/DD neurons (Chase et al., 2004) in light of our observation that optogenetic dopamine neuron activation fails to elevate egg-laying in GABA-deficient mutants (Figure 6A).

How does dopamine influence egg-laying rates during the roaming state, but not the dwelling state? One possibility is that dopamine release dynamics differ in roaming, as compared to dwelling. Alternatively, downstream targets could integrate their detection of dopamine with a locomotion state variable, such that dopamine only enhances egg-laying during roaming. Our results argue against the latter, since elevating dopaminergic neuron activity during dwelling is sufficient to increase egg-laying rates. Moreover, our *in vivo* calcium imaging experiments revealed increased calcium fluctuations in the dopaminergic PDE neurons during high-speed roaming. Specifically, we observed that PDE calcium levels oscillate during forward movement, peaking during a stereotyped phase of the forward propagating bend that overlaps with and shortly precedes the peak phase of egg-laying during roaming. We have not yet determined the underlying mechanism that drives these changes in PDE activity. However, since PDE is a ciliated sensory neuron that expresses mechanoreceptors like *trp-4* (Kang et al., 2010; Li et al., 2011; Sawin et al., 2000), it is possible that PDE might be activated by the animal’s own movement or by the increased flow of external food along the body during high-velocity roaming. In favor of the latter possibility, we found that PDE dynamics were reduced in the absence of food. Thus, it is possible that PDE receives environmental feedback that indicates the degree of movement along a food source. This type of environmental feedback is known to occur for other sensory modalities – for example, optic flow provides animals with visual feedback of how they are progressing through a visual scene.

The egg-laying circuit consists of the HSN command neuron that provides strong input to vulval muscle, as well as VC, VB, and VD neurons that also innervate the vulval muscles. We mapped out downstream effectors of dopamine by optogenetically activating dopaminergic neurons in mutant backgrounds lacking candidate neurons and neurotransmitters. These experiments showed that HSN and VC neurons are important for dopamine’s effects on egg-laying and that GABAergic signaling in VD/DD neurons is required for these effects. Recent studies have shown that GABA receptors are found on a multitude of *C. elegans* neurons that do not receive direct synaptic inputs from GABAergic neurons, including HSN and VCs (Yemini and Hobert, 2019). Given that VD/DD neurons have GABA release sites in close physical proximity to the egg-laying circuit, it is possible that GABA acts extrasynaptically to influence egg-laying.

It is widely appreciated that neuromodulation contributes to behavioral state control, but the widespread behavioral changes that occur across states make it challenging to understand each neuromodulatory system’s specific contribution. The systematic approach described here reveals that in *C. elegans* dopamine elevates egg-laying rates during the roaming state. Previous work has shown that PDF neuropeptides drive high-speed locomotion typical of the state, and have a more modest effect on egg-laying. The combined actions of these neuromodulators and others likely give rise to the full set of behavioral parameters that define the state, though the nature of their interactions will require additional work. Recent studies of mammalian anxiety states have shown that individual neuromodulators like Tac2 neuropeptides can themselves also function in parallel circuits to control different behaviors (Zelikowsky et al., 2018). Studies of fixed action patterns in invertebrates have also revealed parallel functions of single neuropeptides in distinct circuits (Kim et al., 2006; Scheller et al., 1982). The whole-organism behavioral profiling approach that we describe here allows the behavioral changes that differ across states to be more fully characterized, as opposed to being analyzed one behavior at a time. Such a level of understanding will be essential to reveal the neural mechanisms that underlie the generation of these brain-wide states.

## MATERIALS AND METHODS

### Strain List

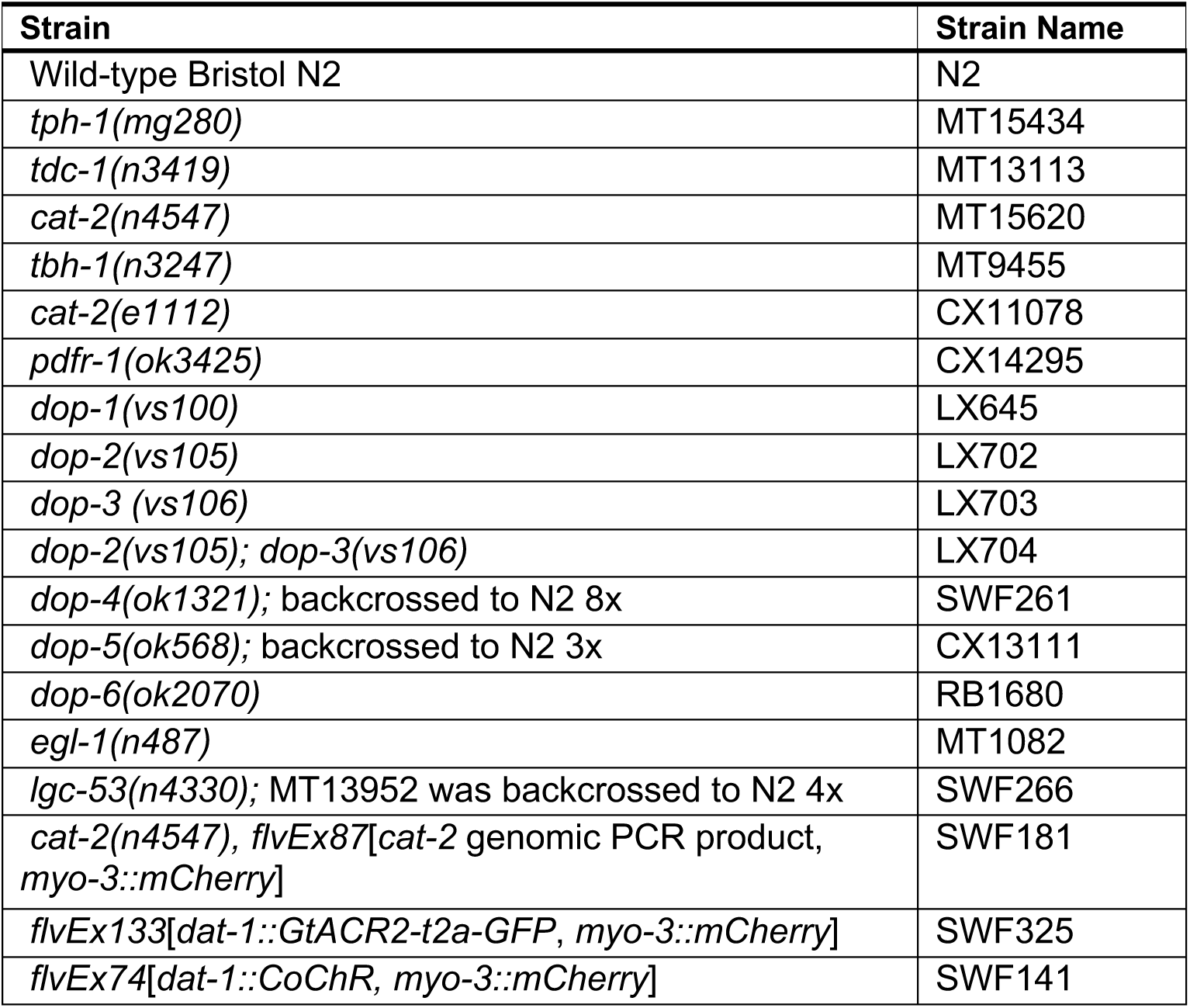
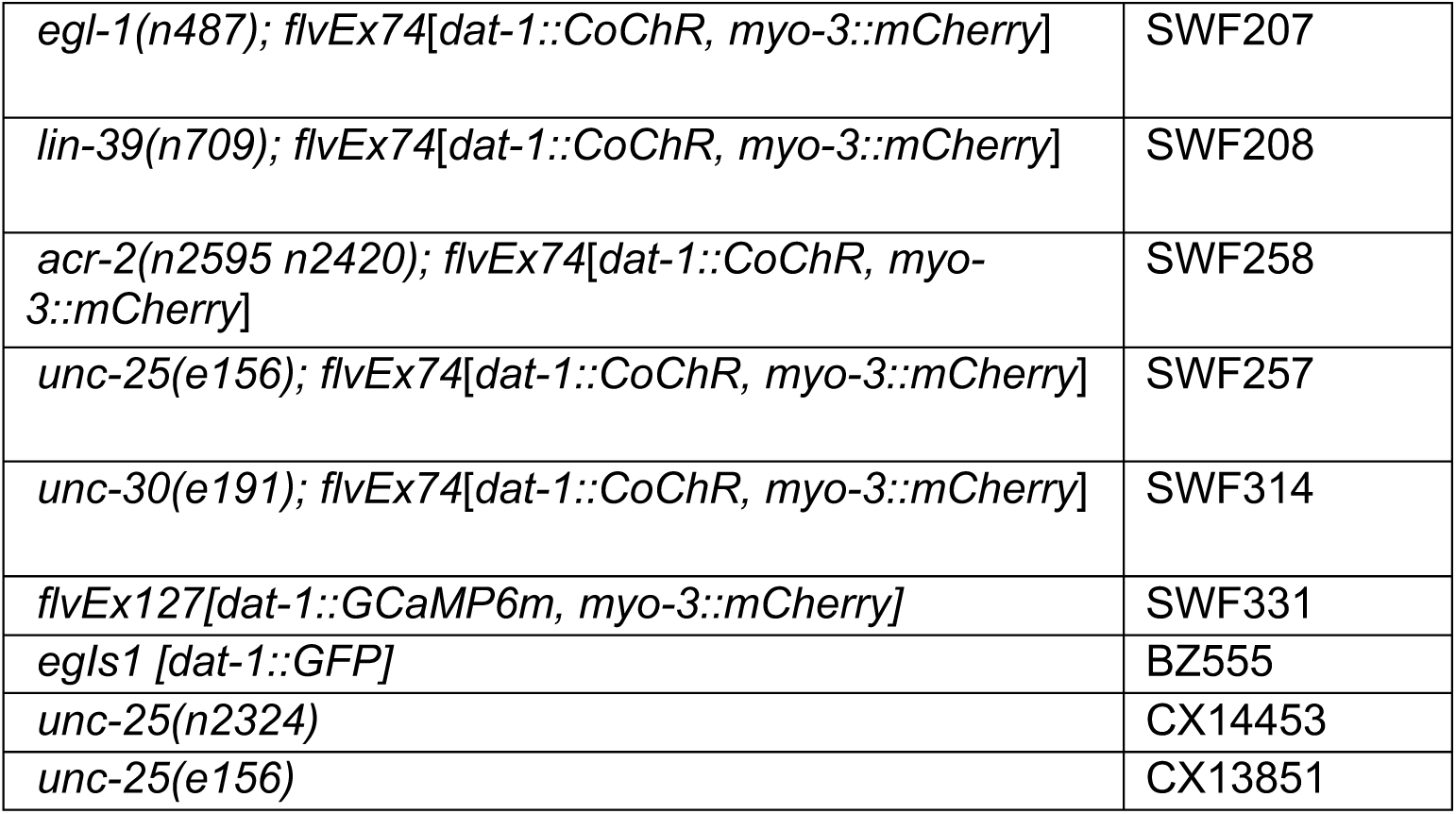

### Growth Conditions and Handling

Nematode culture was conducted using standard methods (Brenner, 1974). Populations were maintained on NGM agar plates supplemented with *E. coli* OP50 bacteria. Wild-type was *C. elegans* Bristol strain N2. For genetic crosses, all genotypes were confirmed using PCR. Transgenic animals were generated by injecting DNA clones plus fluorescent co-injection marker into gonads of young adult hermaphrodites. All assays were conducted at room temperature (∼22°C).

### Data Availability

Data files are publicly available at Dryad (doi:10.5061/dryad.t4b8gthzf).

### Behavioral Recording Conditions

Animals were staged approximately 72 hours prior to recording. 10 adult animals were picked to an NGM plate seeded with *E. coli* OP50 and left to lay eggs for approximately one hour. Adult animals were removed, and eggs were allowed to develop into adults at room temperature until recorded. For experiments that did not involve optogenetics, animals were left undisturbed for the entire developmental period. For experiments that included optogenetics, animals (including controls) were picked as L4s to an NGM plate seeded with a bacterial lawn with 50 µM all-trans-retinol (ATR).

The assay plates were low-peptone (0.2 g/L) NGM plates seeded with 200 µL OP50. 50 µM ATR was added for recordings involving optogenetic stimulation. Roughly circular lawn shapes were created with a spreader, and plates were left to dry overnight at room temperature.

For each on-food recording, a single 72-hour old adult animal was picked to a plate. Plates were taped onto the microscope stage open and face-down on top of 3 evenly placed spacers to prevent condensation on the stage. For animals recorded in the absence of food, a copper ring made of filter paper dipped in 0.02M copper chloride solution was positioned on the plate to prevent the animal from reaching the plate edge. For these recordings, animals were washed in M9 twice and placed on the no-food plate using a glass pipette. Kim wipes were used to remove any remaining liquid.

Recordings began 10 minutes after animals were positioned on the assay plate and were approximately six hours long for on-food recordings and 1-2 hours long for off-food recordings. Optogenetic stimulation was performed with a 532 nm laser at an intensity of 150 µW/mm^2^ for three minutes every ten minutes.

All data were included in analyses, except for a small number of datasets that had very poor quality. The criteria for exclusion, which were uniformly applied, were: (1) if the posture could not be determined in >10% of the frames, or (2) if the head vs tail could not be confidently assigned.

### Tracking Microscope

#### Overview

Recordings were made on three identical tracking microscopes. These microscopes that generate near-comprehensive records of *C. elegans* behavior were named after all-seeing figures: Santa Claus (“he knows when you’ve been naughty or nice”), Sauron (“the Great Eye is ever watchful”), and Cassandra (“all that you foretell seems true to us”). The tracking microscopes and accompanying software suite were inspired by and share similarities to previous imaging platforms (Yemini et al., 2013). Details of microscope design and construction are provided here. The complete bill of materials and build instructions for the tracking microscope are available in a git repository at https://bitbucket.org/natecermak/openautoscope. The image analysis code in R is also available at https://bitbucket.org/natecermak/wormimageanalysisr.

#### Microscope optics

The tracking microscope was built predominantly using optics and an optical cage system from Thorlabs. The basic optical path consists of a 4x NA 0.1 Olympus PLAN objective (Thorlabs #RMS4X) with a 150 mm tube lens (Thorlabs #AC254-150-A-ML), yielding a magnification of 3.33X. The image is collected on a monochrome Chameleon3 camera (FLIR, #CM3-U3-13Y3M), with a 1/2” sensor format and 1280×1021 pixels (4.8 μm/px at sensor plane, 1.44 μm/px at object plane). This yields a field of view of 1.84 x 1.47 mm. We illuminated the sample with a 780 nm, 18 mW IR LED (Thorlabs #LED780E) in a transillumination configuration. The LED was diffused using a ground glass diffuser (Thorlabs #DG10-120), and collimated onto the sample using a 16 mm aspheric condenser lens (Thorlabs #ACL25416U) placed 16 mm from the diffuser surface.

The microscope path also included a 4.5 mW, 532 nm laser diode module (Thorlabs #CPS532) for optogenetic stimulation. The laser beam (3.5 mm diameter) was combined with the main optical path using a 550 nm dichroic (Thorlabs #DMLP550R), then focused on the back focal point of the objective using a f=100 mm planoconvex lens (Thorlabs #LA1509-A-ML). This illuminated a circular region roughly 1.6 mm in the sample plane, with a calculated average intensity of 150 μW/mm^2^.

#### Microscope mechanics

We reasoned that mechanical motion of the stage might perturb the worm, so we designed our tracking microscope to move the optical components rather than the sample. We attached the optical train to a gantry on two motorized linear actuators, enabling translation of the optics in the X-Y plane (perpendicular to the optical axis). However, we moved the sample in the Z-axis for focusing.

We used motorized C-beam linear actuators from OpenBuilds Parts Store, which allow roughly 150 mm of travel distance in each axis. They use an 8 mm-pitch leadscrew to translate rotational into linear motion. We drove these leadscrews with NEMA23 stepper motors (Pololu, #1476). Because these motors have 200 steps per revolution and we drove them with 1/32 microstepping, the single-step resolution was 1.25 μm in each linear axis.

To control the motors, we used DRV8825 stepper controller ICs. These are common stepper motor driver ICs used in CNC and 3D-printing applications. The DRV8825 ICs were controlled by a Teensy 3.2 microcontroller with custom-written firmware, using the AccelStepper library for smooth stepping control. The Teensy 3.2 firmware communicates with the computer over a high-speed serial connection via USB, and continually broadcasts its current position at 200 Hz. The computer controls movement by sending strings to the Teensy telling it to move to a given absolute coordinate – for example “X200” means go to position 200 in the X-axis, whereas “Z-3000” means go to position −3000 in the Z-axis. Each axis was programmed to move at a maximum speed of 1000 microsteps/s (1.25 mm/s), with a maximum acceleration of 50,000 microsteps/s^2^ (62.5 mm/s^2^) for X and Y, and 5,000 microsteps/s^2^ (6.25 mm/s^2^) for Z.

#### Tracking

Our tracking approach was similar to previous reports (Stephens et al., 2008). We wrote a custom LabView program (National Instruments) to perform tracking. In this program, we acquired and processed images at ∼20 Hz. For each frame, we thresholded the image, dilated and eroded the binary image to smooth the outline and fill small holes, and detected connected components. Connected components outside a given size range (typically 20,000-100,000 pixels) were discarded, and of the remaining components, the one nearest to the center of the frame was selected as the target. We calculated the unweighted centroid of that component, and used its deviation from the center of the frame as the error signal. We used a proportional-integral (PI) controller to ensure the worm stayed centered in the image. Typical P and I gains were 0.3 steps/pixel and 0.1 steps/(pixel*second), respectively. The feedback loop rate was set by the frame rate (20 Hz). While we could track worms equally well at faster frame rates (easily exceeding 40 Hz), we found 20 Hz both provided stable and robust tracking. Additionally, 20 Hz was near optimal for recording because below 20 Hz we were unable to accurately quantify pharyngeal pumping, and sampling faster than 20 Hz provided essentially no additional information about worm behavior.

During tracking, each frame was saved as a jpeg file. To reduce the file size, we did not save the entire frame, but instead cropped the image to the target component (described above), padded by 40 pixels in all directions. Typical frames were on the order of 20 kB after jpeg compression.

Because the agar surface is not perfectly level, the microscope must periodically refocus as the worm moves. To automatically focus, we performed a rapid z-series centered on the current z-plane and calculate the sharpness of the image at each z-position. Sharpness was calculated as the variance of the image after high-pass filtering with a Laplacian filter. The microscope then returned to the z-plane that yielded the sharpest image. The entire autofocus operation took ∼2.5 seconds. Autofocusing was triggered 10 minutes after the previous autofocus, and every time the worm moved more than 2.5 mm from the position of the last autofocus.

In our experience, tracking was extremely robust and worms never exceeded the boundary of the camera frame under typical conditions. However, there were two cases in which tracking sometimes failed – when worms reached the boundary of the dish and when condensation on the plate lid reduced contrast or overall image brightness. When worms reached the dish boundary, there were often dark regions in the image that the worm might enter, at which point the tracking code would no longer be able to identify the worm’s position. Similarly, condensation in the optical path tended to decrease overall brightness and contrast, which in some cases prevented the code from identifying the worm as a dark component on a light background.

### Behavior detection

While gross locomotor behavior could be quantified online, most other metrics were not, due to computational requirements. We developed an offline analysis pipeline in R (R Core Team, 2019) to 1) calculate additional locomotion-related metrics and also quantify 2) body posture 3) pumping 4) defecation and 5) egg-laying.

#### Locomotion

We quantified multiple metrics of locomotion on 1- and 10-second timescales, as these capture different aspects of behavior. We measured speed as the magnitude of the displacement of the worm’s center of mass and velocity as displacement projected onto the worm’s body axis. We also measured the angular direction of the worm and calculated the angular velocity as change in angular direction over time. Finally, we tracked the distance between the worm’s center of mass and the bacterial lawn (equal to 0 if the worm was on the lawn), as well as the distance from the worm’s nose to the lawn.

#### Body posture

Body posture has been the subject of extensive study in *C. elegans* (Stephens et al., 2008; Yemini et al., 2013). We used a well-established pipeline to quantify body posture. This pipeline follows the same protocol as the online tracking analysis (above), including thresholding, dilation and erosion, and selecting the connected component representing the worm. That component is then thinned by iteratively applying a 3×3 look-up-table based thinning filter. This process yields pixels that are along the worm’s centerline. We treat these pixels as a graph, in which bright pixels become nodes and we create edges between all pairs of adjacent (8-connected) pixels. If the graph contains cycles, we mark the frame as “self-intersecting” and do not analyze it further. “Self-intersecting” frames are most commonly due to the animal exhibiting an omega bend, or a related body posture. If the graph has branchpoints, we iteratively prune the shortest branches until the graph no longer has branchpoints and is thus a single line. We refer to this as the centerline. We smooth the centerline coordinates (31-pixel running mean) to reduce discretization artifacts, and resample the centerline so that it contains 1001 evenly-spaced points. We calculate angles between all consecutive points, and fit this vector of angles to a 2nd-degree b-spline basis consisting of 14 components. This yields a 14-dimensional vector that accurately and succinctly represents the worm’s posture. We record the mean and interpret it as the worm’s overall orientation. We mean-center the 14-point vector, which we then interpret as representing deviations from the worm’s overall orientation. Because this representation allows straightforward and accurate reconstruction of the worm’s centerline, we also use this centerline to calculate curvature along the centerline as the inverse of the local radius of curvature.

#### Assignment of head/tail

Our first steps to assign head and tail to each frame was to simply ensure consistency frame-to-frame. The initial body posture quantification described above occurred without regard to which end is the head and which is the tail. However, we then subsequently enforced angular consistency between consecutive frames. To do so, we iterated through frames and if the mean angle difference between frames *i* and *i+1* exceeds π/2, we reversed the orientation of the worm by reversing the angle vector and subtracting π from all the body angles.

This was insufficient to ensure consistency across entire datasets, as some frames did not have body angles assigned due to self-intersection of the centerline (described above). We then iterated through contiguous blocks of non-self-intersecting frames and ensured that each block had the same average centerline intensity (brightness) profile as the previous block. Specifically, we checked whether the average centerline intensity vector of block *j+1* or its reverse was better correlated to that of block *j.* If the reverse was a better match, then the entirety of block *j+1* was reversed.

Finally, having verified that the orientation is at least consistent across the entire experiment, we manually annotated which end was head and which was tail.

#### Pumping

Our general strategy to quantify pumping was to detect the position of the grinder as well as the posterior end of the posterior bulb and then count negative peaks in the distance between them. This approach is similar to Lee et al. (2017), but performed on a freely-moving worm instead of a worm immobilized in a microfluidic chip.

Because there were occasional errors in estimating the worm’s centerline (typically due to the presence of eggs touching the body, especially the tip of the nose/tail), we first eliminated from the analysis all frames in which the worm’s length suddenly increased. We identified these frames as those in which the worm length exceeded the 90^th^ percentile of the surrounding 10 minutes by at least 30 μm.

Within the remaining frames, we identified the grinder and the posterior end of the posterior bulb as dark positions in the intensity (brightness) profile along the worm’s length. This profile was obtained by averaging the intensity of 8 perpendicularly-oriented pixels for each point on the centerline. We then used spline interpolation to upsample this profile to a roughly 0.2 μm resolution, and looked for the presence of two local minima in the profile. In particular, we sought two local minima amongst the four darkest local minima in the first quarter of the worm’s length that were between 8 and 17 μm apart. If these were found, we calculated the distance between them, otherwise we marked the frame as missing data. We refer to this signal as the grinder distance.

For contiguous stretches for which the grinder distance was defined, we bandpass filtered this signal (2nd-order Butterworth filter, passband 0.2-0.8 normalized frequency) and counted peaks that deviated more than 0.6 μm below 0 (the average grinder position). Pumping rates were only defined over regions in which the grinder distance was not missing.

#### Defecation

We used the stereotyped body contraction, known as the defecation motor program (Thomas, 1990), to identify defecation events. We first identified proportional fluctuations in the worm’s body length by subtracting a moving median filtered (width 50 seconds) estimate of the worm’s length, then dividing by the value of the median filter. This yielded a signal with roughly zero mean, in which deviations represented the proportion by which the length changed. We then applied a matched filter with a kernel consisting of two Gaussians (width 0.9 seconds) separated by 3.5 seconds. This is a rough approximation of the body length profile during a typical defecation event. Peaks in the filtered signal were taken to be defecation events.

#### Egg-laying

Our general strategy for detecting egg-laying was to look for sudden increases in the worm’s width around the vulval region.

We calculated the worm’s body width by measuring the distance from the worm’s centerline to the dorsal or ventral edge of the worm, perpendicular to the centerline, at 1001 evenly spaced points along the centerline. To compress these oversampled vectors, we then fit these to a b-spline basis consisting of 30 components, yielding two 30-point “half-width” (centerline to edge) vectors.

We developed a heuristic in which candidate egg-laying events are detected by meeting the following criteria:

1. the worm’s area does not decrease
2. the worm’s length does not change by more than 5%
3. the worm’s width in any of segments 15-20 (roughly the midpoint to 66% along the worm’s length) on either side increases abruptly within 100 ms by at least 6 pixels.
4. a filter calculated using the worm’s width crosses a threshold. In particular, this filter was designed to maximize the signal from an egg suddenly appearing around the vulva, while minimizing the signal from a worm moving alongside an existing stationary egg.

The latter filter was constructed using the average width of 5 segments anterior to the vulval region, 5 segments around the vulval region, and 5 segments posterior to the vulval region. A differencing filter of length 250 ms was applied to the width time series from each 5-segment region to detect arrival of eggs, and then the three regions were summed with weights 1, 2, and 1 respectively.

Because *C. elegans* only lays eggs on its ventral side, our analysis then identified the putative ventral side as the side of the worm with more candidate egg laying events. This criterion proved highly reliable in that it correctly identified the ventral side for every worm we analyzed (verified during manual annotation, below). Candidate egg laying events on the dorsal side were ignored.

These criteria yielded generally high sensitivity but poor specificity, often identifying up to >200 frames containing candidate egg laying events (out of >400,000 frames for a six-hour dataset). We then manually verified these candidate frames, typically identifying 20-70 real egg-laying events. For each event we also manually identified the number of eggs laid, as multiple eggs are occasionally laid simultaneously.

#### Lawn boundary

Before each experiment, we manually annotated the position of the bacterial lawn by manually directing the microscope to travel around the perimeter of the lawn and record the path coordinates. This enabled post-hoc determination of when the worm was on the bacterial lawn and if not, how far the worm was from it.

### Posture Analysis and HMMs

#### Overview

As is described in the main text, we used time-varying changes in *C. elegans* posture to learn behavioral states displayed by adult animals feeding on OP50. This pipeline consisted of several data processing steps, described here.

#### Data pre-processing

The input data for these analyses consisted of 14-element data vectors at each time-point that describe the worm’s posture as a series of relative body angles, from nose to tail (see “Behavior Detection” section above). We used the location of egg expulsion along the body to determine the ventral side of the body, and inverted the body angles of animals travelling on their left sides, so that all animals were aligned along the same dorsal-ventral axis.

#### Posture Compendium

To express the worm’s posture in a simple, compressed manner, we constructed a compendium of reference postures that encompass the broad range of postures that *C. elegans* animals display while feeding. This approach has been previously employed by other groups (Schwarz et al., 2015). Then, for each posture observed at each moment in time, we determined the closest match in the compendium using k-nearest neighbors. To construct a compendium, we sampled 100,000 frames from 24 N2 animals and performed hierarchical clustering. We then cut the dendrogram at different depths (1-200) to build compendiums with different numbers of reference postures. Each time, we averaged together the body angles in each cluster to obtain the prototypical posture for that cluster. We could describe the degree to which the compendium faithfully captured the worm’s posture by measuring how much of the variance of the body angle vector was discarded when transforming the actual posture vector to its most similar reference posture. Then, to determine the optimal size of the posture compendium, we examined the variance explained for compendiums with a wide range of sizes (1-200). This curve (Figure 2—figure supplement 1A) clearly plateaus beyond a compendium size of ∼70, indicating that adding additional postures to the compendium beyond this point fails to capture a great deal more variance. We identified the size at which at least 75% of the postural variance was explained while adding another posture explained less than 1% of additional variance. This size was 100 postures, and the 100-posture compendium explained approximately 76% of the variance (Figure 2— figure supplement 1A).

#### Transitions between reference postures

To begin identifying stereotyped postural changes, we examined the probability of each reference posture transitioning to the other reference postures. In this analysis, we ignored self-transitions. This transition matrix was constructed using 6 animals. We then clustered the rows of this transition matrix using k-means clustering to identify sets of postures with similar transition profiles. We observed a clear, symmetric, block-like structure to the clustered transition matrix, indicating that there are sets of postures that animals tend to transition back and forth between. We performed k-means clustering 500 times for a given number of clusters, while varying the number of clusters from 2-10. To determine the optimal number of clusters, we measured the degree to which the block-like structure emerged for a given clustering by calculating the ratio of the average intra-group transition probabilities and the average inter-group transition probabilities. Based on this measurement, the optimal number of clusters was found to be eight, and we selected the clustering with the highest value for further analysis. We refer to these eight clusters of reference postures as “posture groups.”

#### Pre-processing for HMMs

To capture slow time-scale changes in body posture, we attempted to describe the typical postures emitted by *C. elegans* animals over three-second intervals (60 frames). We chose this time frame because we found that over three-second intervals animals typically generated postures from a single posture group, suggesting that this time frame was a good match to the posture group transition rate. Nevertheless, because animals sometimes generated postures from two or three groups within a give 3s bin, we segmented these three-second intervals by clustering. To perform clustering, we described the posture groups emitted over each three-second interval as an eight-element vector where each entry is the fraction of data points belonging to each posture group. We then clustered these vectors by k-means clustering and used silhouette criterion to determine the optimal number of clusters. The optimal number was found to be eight; there was a clear mapping where each cluster consisted of data vectors primarily belonging to a single posture group. This allowed us to describe each three-second interval with a single value 1-8, based on which cluster it mapped to. We will refer to these cluster values as “coarsened posture groups” in the section below.

#### Training and evaluating HMMs

We trained HMMs on the sequences of coarsened posture groups from 6 animals. The transition and emission matrices were randomly initialized for each model training. To determine the optimal number of hidden states, we trained models across of a range of hidden states (2-12) and used the Bayesian information criterion (BIC) to compare model likelihoods to one another. The curve of median BIC values demonstrated an optimal value at nine hidden states. We trained 200 nine-state models, each from random initial conditions; half were trained on the same group of 6 animals, while the other half were trained on a separate, non-overlapping group of 6 animals. We then evaluated the similarity of these 200 models by measuring the average Euclidean distance between the rows of the emission matrices, i.e. how similar the emissions were in each state. We found that the models with the highest likelihoods were essentially identical, despite being trained from different sets of animals and different initial conditions (Figure 2—figure supplement 1E-F). This observation suggests that model training reliably converges to the same solution. We arbitrarily selected one of these extremely similar models for further use. To determine the most likely state path for each animal, we used the Viterbi algorithm.

#### *In vivo* calcium imaging

*In vivo* calcium imaging of the four dopaminergic cell types was conducted on one-day-old wild-type animals. Animals were positioned on thin, flat slices of NGM agar with a PDMS outer barrier so that animals could not crawl off the agar. OP50 *E. coli* was seeded uniformly on the agar, except for off-food videos. Animals were permitted to equilibrate on the agar slides for 10 min prior to the beginning of each recording. Animals were then recorded with a 4x/0.2NA Nikon objective and an Andor Zyla 4.2 Plus sCMOS camera. Blue light output to animals was 9-20% output from an X-Cite 120LED system for 10 msec of each exposure. Brightfield images (to collect postural and behavioral information) and fluorescence images were collected in an alternating fashion, made possible by alternating which light source was active during each camera exposure. The total frame rate was 20fps, so that each channel (GCaMP and brightfield) had an effective frame rate of 10fps. GCaMP data and behavioral data were thus collected with a 50ms lag, though for the purposes of our analyses we dis-regarded this small time lag. GCaMP fluorescence was tracked using a previously-described ImageJ tracking macro (Flavell et al., 2013) and behavioral data were extracted using the software that was developed for the tracking microscope analysis (described above).

## Author Contributions

N.C., S.K.Y., R.C., Y-C.H. and S.W.F. designed experiments. N.C., S.K.Y., R.C., Y-C.H. and S.W.F. conducted experiments, analyzed data, and interpreted data. N.C., S.K.Y., R.C., and S.W.F. wrote the paper.

## Acknowledgments

We thank Cori Bargmann and members of the Flavell lab for helpful comments on the manuscript, Joshua Powers and Karen Gao for assistance with data annotation, and Andrew Bahle for help with microscope construction. We thank the Bargmann lab, Horvitz lab, and the *Caenorhabditis* Genetics Center (supported by P40 OD010440) for strains. Y-C.H. acknowledges support from the Picower Fellows program. S.W.F. acknowledges funding from the JPB Foundation, PIIF, PNDRF, the NARSAD Young Investigator Award Program, NIH (R01NS104892) and NSF (IOS 1845663 and DUE 1734870).

## Supplementary Legends

**Video S1. Examples of behavioral states captured through posture-HMM.** This video shows examples of the nine different behavioral states identified through posture-HMM. Note that videos are of different durations and are looped.

**Figure 1 – Figure Supplement 1.**
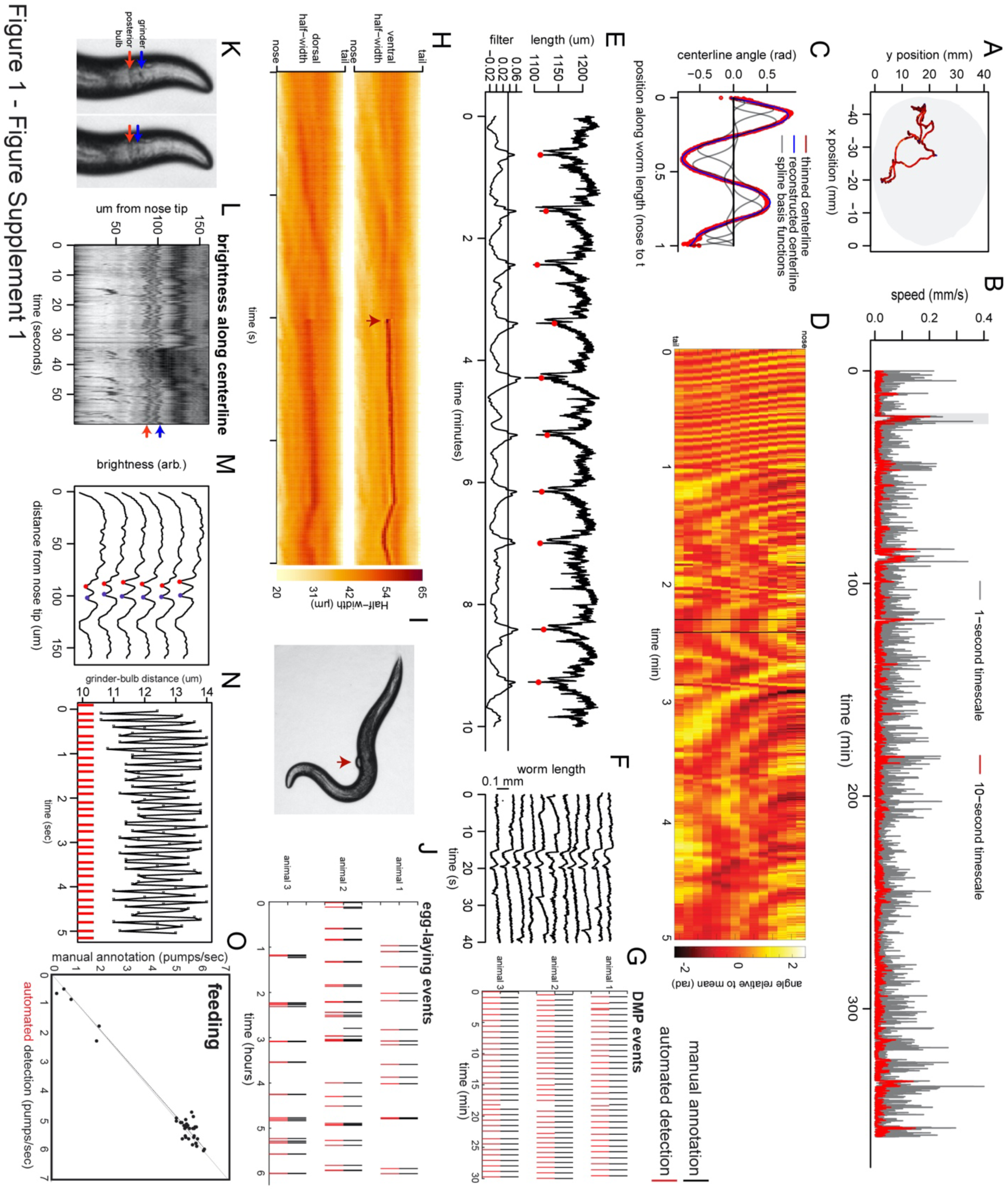
Extraction of behavioral parameters from video recordings, and validation of methods. (A) Movement path of *C. elegans* animal during recording on tracking microscope. Path is color-coded by velocity (black is slow; red is fast; blue is reverse). (B) Example speed traces over a 6-hour recording of a wild-type animal. Gray shows speed calculated with a 1-second time step; red shows it calculated with a 10-second time step. (C) Body angles are accurately captured by 14-point parameterization. Data are body angles from a worm in one example frame. Centerline angles are compressed to a 14-point vector consisting of weights for a spline basis (basis functions shown as grey lines). (D) Body angles over a five-minute example dataset. Missing data (due to self-intersecting loops in body posture) are in black. (E) The defecation motor program (DMP) is detectable as a stereotyped pattern of body length contractions. Top line shows body length over 10 minutes. Bottom shows the data after filtering with a kernel approximating the shape of the DMP body contraction. The filter enables straightforward thresholding to detect defecation events (red dots). (F) Ten sequential defecation events are shown, revealing the stereotyped nature of body contractions during defecation events. (G) Validation of DMP detection. For three animals recorded on different microscopes, we compared the automated detection of DMP events to manual user annotation for a random 30-minute segment of each video. Rasters indicate detected DMPs. (H) Egg-laying events are detectable via instantaneous increases in body width. Example of an egg-laying event, with the widths of the dorsal and ventral sides of the body shown. Note that the width increases at the mid-body suddenly (arrowhead). This increase is limited to the ventral side, where the vulval muscles are located. (I) Image of a worm at the moment of an egg-laying event for the dataset in (H). (J) Validation of egg-laying detection. For three animals recorded on different microscopes, we compared the automated detection of egg-laying events to manual user annotation for all six hours of each video. Rasters indicate detected egg-laying events. (K) Pharyngeal pumping is detected via movement of the grinder. Two sequential frames (50 ms apart) illustrating pharyngeal pumping. Pumping involves rapid retraction of the grinder, relative to the posterior end of the terminal bulb. (L) The grinder and posterior end of the terminal bulb can be detected as local minima in centerline brightness profiles. Heatmap shows brightness along the centerline for the first 160 μm of the worm’s length. Red and blue arrows show the positions of the grinder and posterior end of the terminal bulb, respectively. (M) Six sequential frames of the data in (L), plotted as line traces. Traces are offset for clarity. Red and blue dots denote the detected position of the grinder and the posterior end of the terminal bulb. (N) Pumping is clearly visible as negative peaks in the distance between the grinder and the posterior end of the terminal bulb. Plot is a time series of the distance between the grinder and posterior end of the terminal bulb, showing oscillations (pumps) at roughly 5 Hz. Red ticks at the bottom denote detected pumps. (O) Validation of pumping detection. For 35 short (20 sec.) video segments (taken from 15 different recordings), we compared the automated detection of pumping rates to manual annotation by users watching videos in slow motion. R^2^ value is 0.9671.

**Figure 1 – Figure Supplement 2.**
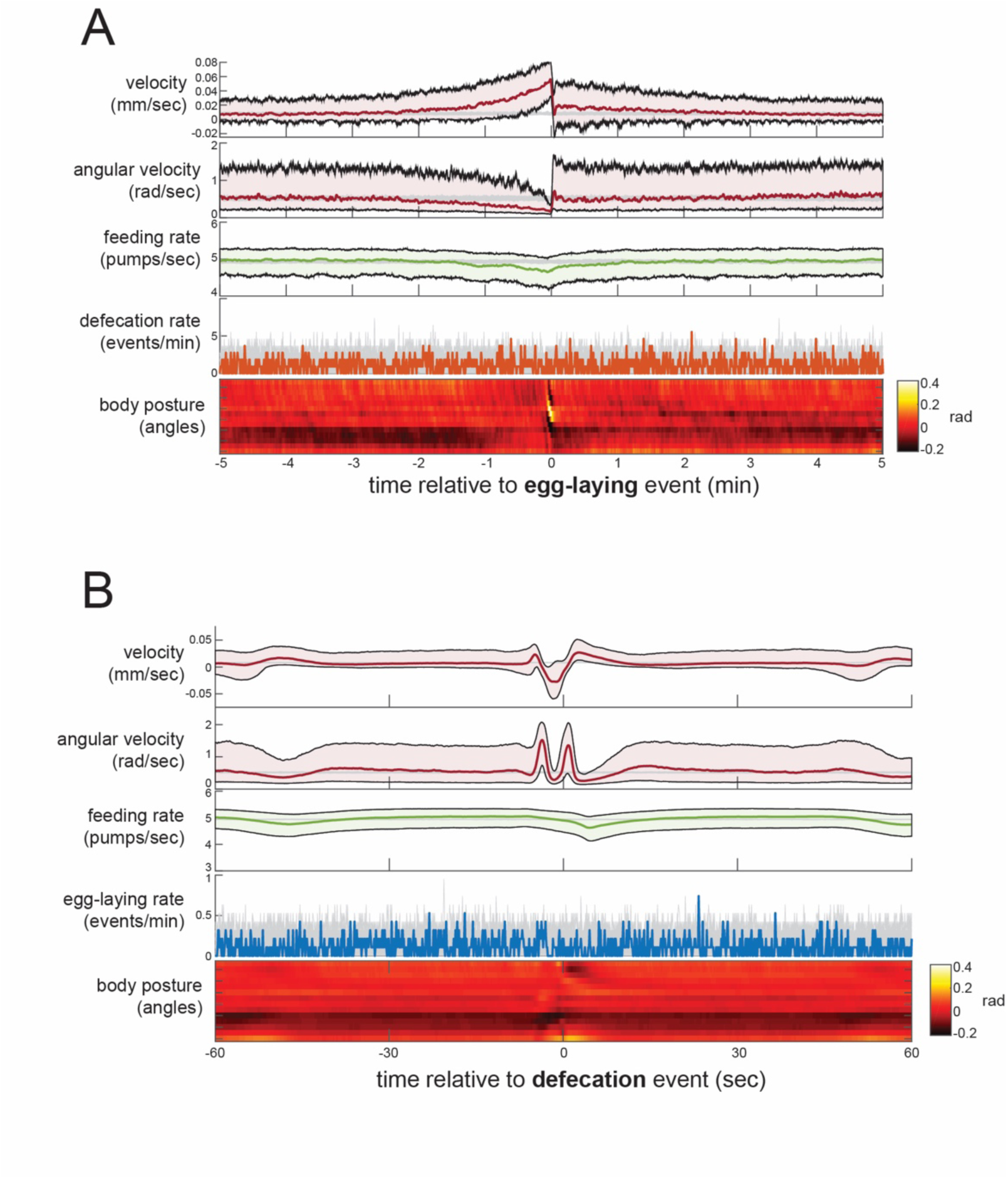
Event-triggered averages showing behavioral coordination surrounding egg-laying and DMP events. (A) Event-triggered behavioral averages surrounding egg-laying events. Velocity, angular velocity, feeding rate, and defecation rate are shown as medians ± 25^th^ and 75^th^ percentiles (gray lines indicate random samples of identical size). Average body posture is shown as a heatmap of angles along the body. (B) Event-triggered behavioral averages surrounding DMP events, shown as in (A).

**Figure 2 – Figure Supplement 1.**
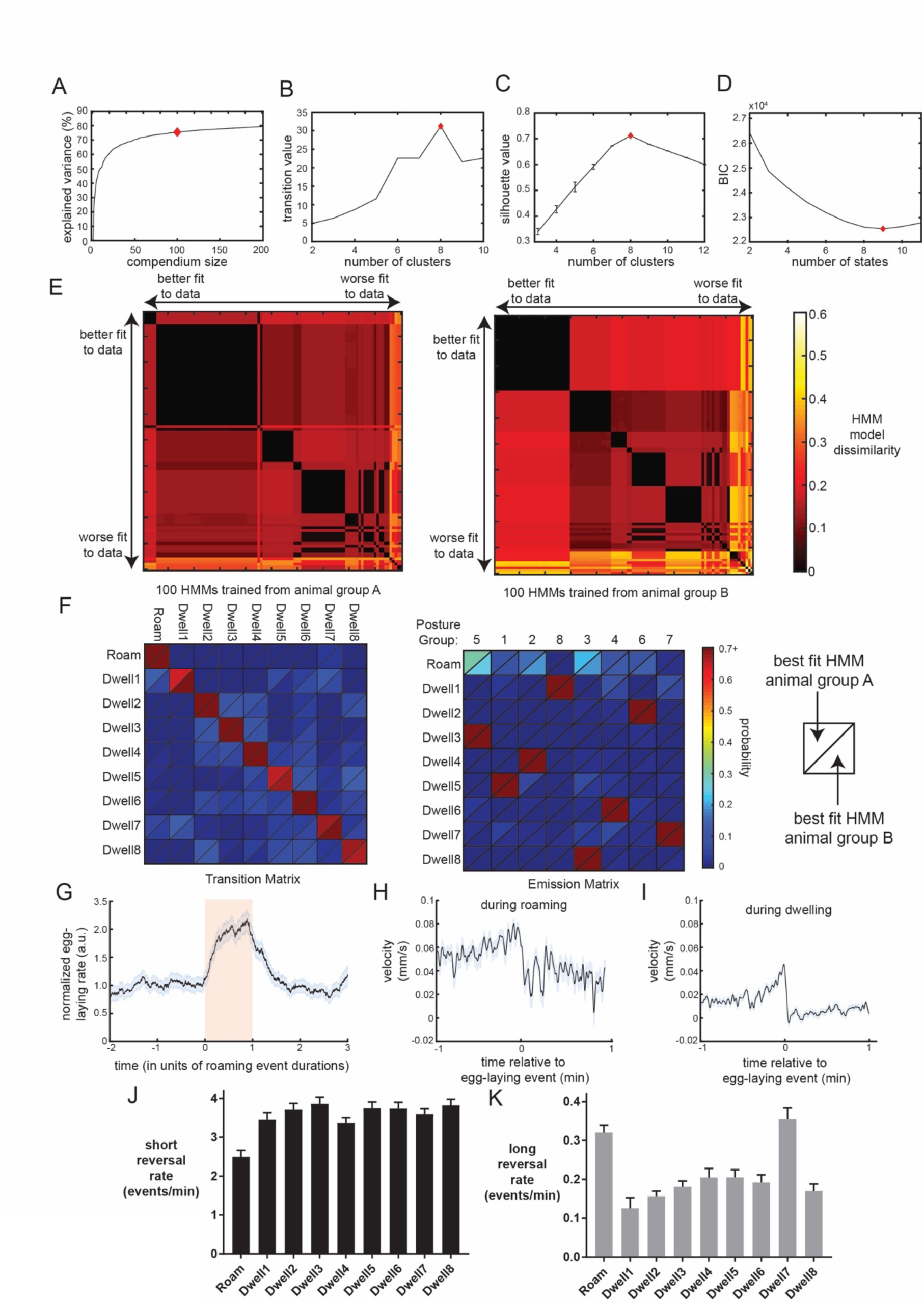
Additional data related to Posture-HMM. (A) A compendium of reference postures can explain most of the variance of observed animal postures. Increasing the number of postures in the compendium leads to a higher percentage of variance explained, though the impact of adding more postures diminishes gradually (i.e. plateaus). Red diamond indicates the number of reference postures used in all related analyses in this study. (B) Clustering of the transition matrix between discrete body postures maximally separates posture groups when 8 clusters (red diamond) are used. “Transition value” is the average intra-group (i.e. intra-cluster) transition rate divided by the average inter-group transition rate. Based on this criterion, the optimal number of clusters was eight. (C) Silhouette value when clustering the 3-second binned posture data in preparation for HMM training. Based on this criterion, the optimal number of clusters was eight. (D) BIC value for HMMs trained with varying numbers of hidden states. Based on this criterion, the best fit to the data was with a nine-state HMM. (E) Similarity of trained HMMs, shown for 100 different HMMs trained on two separate groups of animals. Training for each separate HMM was initialized with random starting parameters. Heat map shows HMM model dissimilarity, which is the average Euclidean distance between the rows of the emission matrices for the two HMMs being compared. HMMs are ordered based on their fit to the full set of wild-type data. Note that the best fit HMMs are very similar to one another (black color). (F) Comparison of the best fit HMMs from animal group A and animal group B shows that the training converged to a very similar solution, despite having totally independent training datasets and different random initial conditions. Data are displayed as heatmaps depicting the transition and emission matrices of the two HMMs. HMM parameters from both states are depicted in each quadrant, separated by a diagonal line (as depicted). (G) Egg-laying rates over the durations of roaming states. All roaming states were “stretched” so that they could be properly aligned, and then the normalized egg-laying rates over the durations of the states were plotted. Note that egg-laying rate are high during most of the roaming states. However, rates are not as high at the very beginnings of these state and the rates are still high for a short while after roaming states terminate. Data are from 1573 roaming states across 30 animals. (H) Event-triggered average of velocity, aligned to egg-laying events. This panel only shows data surrounding egg-laying events that were embedded in roaming states (animal roamed from −1 min to +1min). (I) Event-triggered average of velocity, aligned to egg-laying events. This panel only shows data surrounding egg-laying events that were during dwelling states. (J) The rate of short reversals/reorientations (<4 sec) varies across the Posture-HMM states. (K) The rate of long reversals/reorientations (>4 sec) varies across the Posture-HMM states. For (J-K), n=30 animals.

**Figure 2 – Figure Supplement 2.**
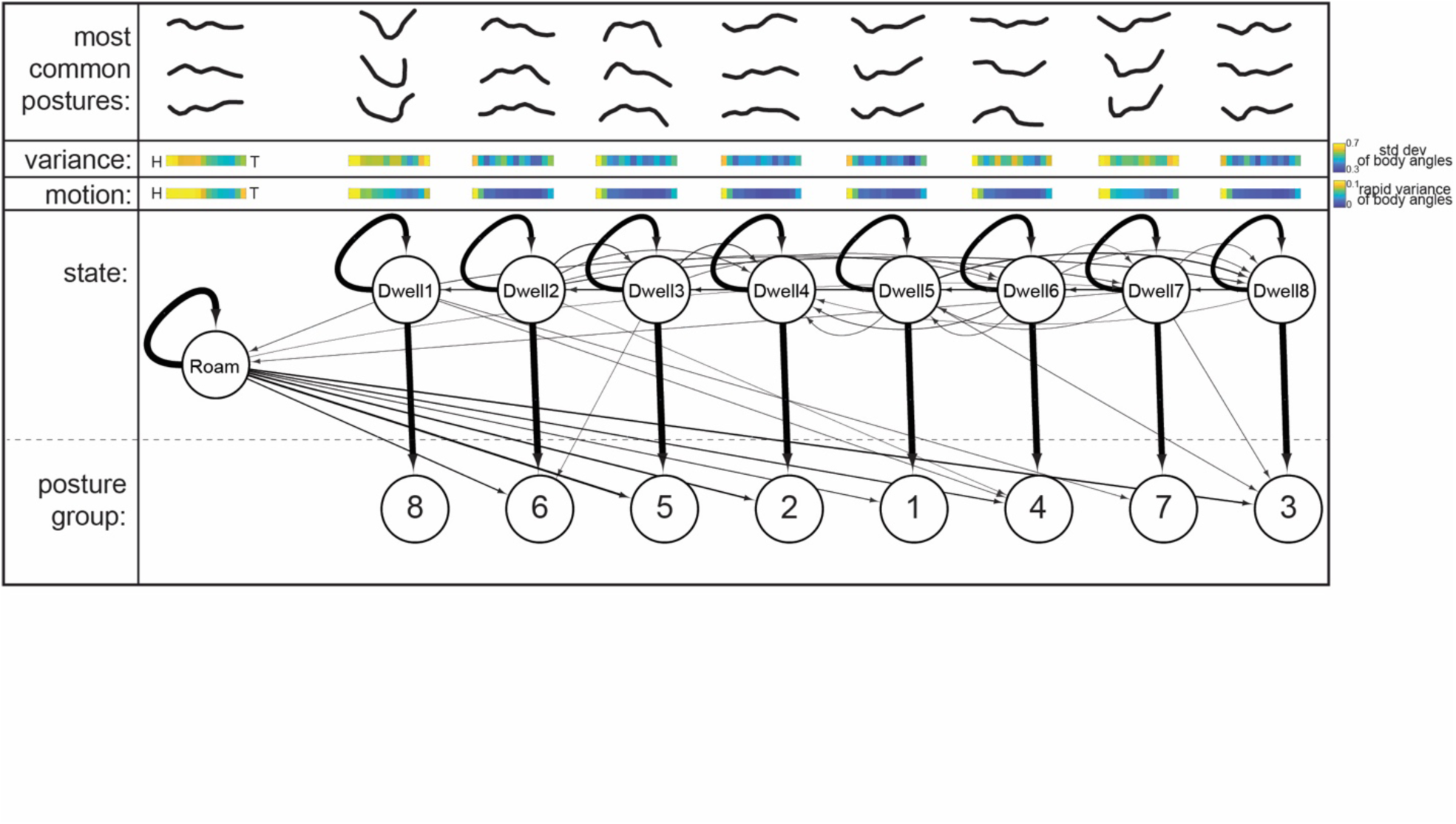
HMM to characterize behavioral states based on postural changes. Bottom: HMM structure learned from behavioral recordings. Arrows between states depict transition probabilities, while arrows directed to posture groups depict emission probabilities. Arrow thickness indicate probabilities. Above: for each state, the most common postures observed while animals are in the state are shown. The postures exhibited in each of the nine states are significantly different from those exhibited in all other states (p<0.05, Bonferroni-corrected permutation test). In addition, “variance” shows the standard deviation of body angles from head to tail across all data points where animals were in the state, providing a measure of how much the body angles vary within the states. “Motion” shows the average variance of body angles from head to tail over short (2 sec.) time intervals, providing a measure of how much rapid-timescale motion is observed at each body angle within each state.

**Figure 3 – Figure Supplement 1.**
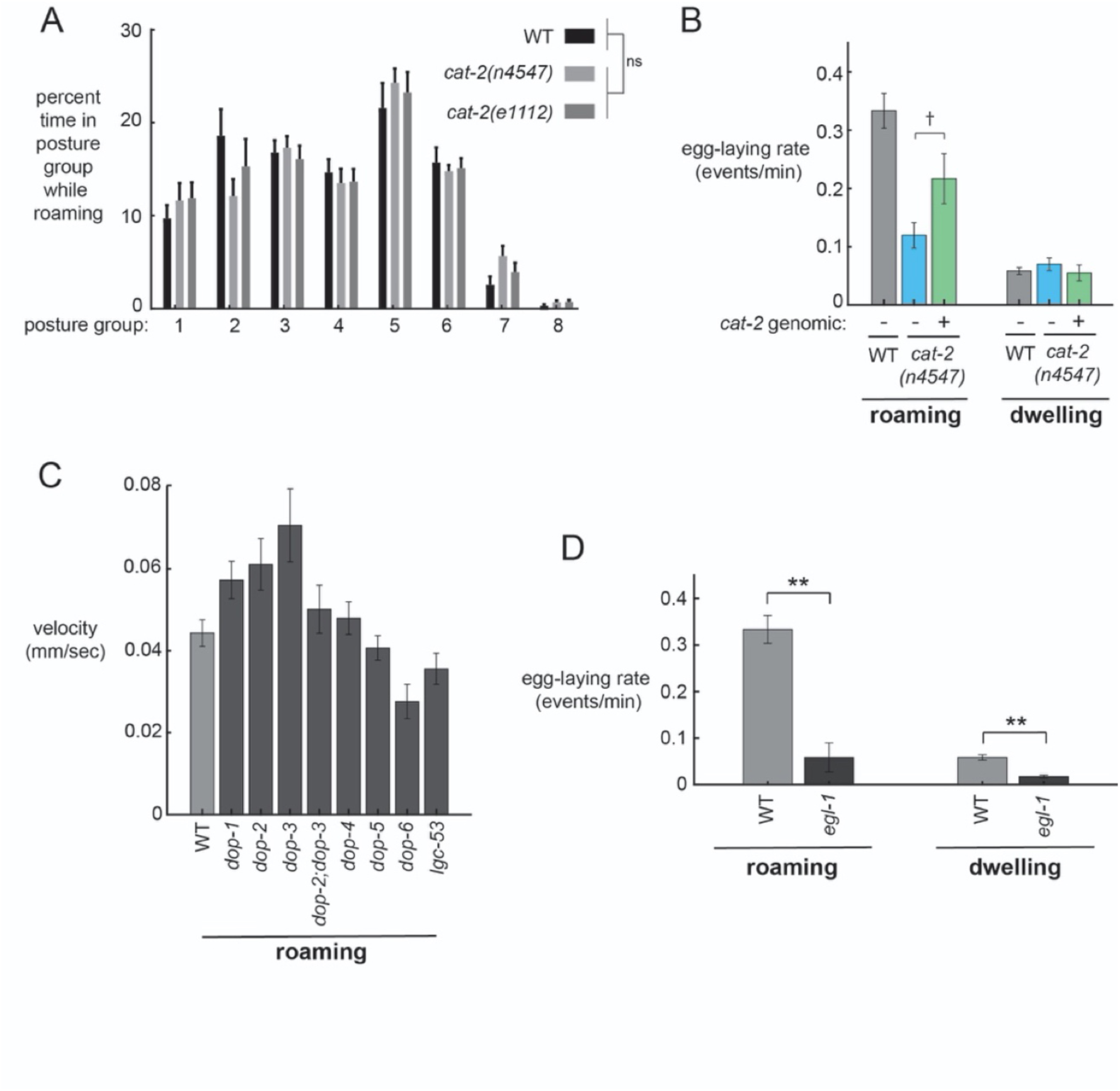
Additional behavioral analysis of mutant strains. (A) The postures that *cat-2* mutant animals display during roaming are similar to those displayed by wild-type. Data are histograms of percent time in each of the eight posture groups during roaming for the indicated genotypes. ANOVA shows no effect of genotype on posture group prevalence. (B) Introduction of *cat-2* genomic fragment into *cat-2(n4547)* mutants elevates egg-laying rates during roaming. n=10-30 animals per genotype. †p=0.0779, t-test. (C) Velocity of mutant animals lacking specific dopamine receptors. Data are means ± standard error of the mean (SEM). n = 9-30 animals per genotype. (D) Egg-laying rates during roaming and dwelling for *egl-1* mutant animals. Data are means ± standard error of the mean (SEM). **p < 0.001, Bonferroni-corrected Welch’s t-test. n=21 animals for *egl-1* and n=30 animals for WT.

**Figure 4 – Figure Supplement 1.**
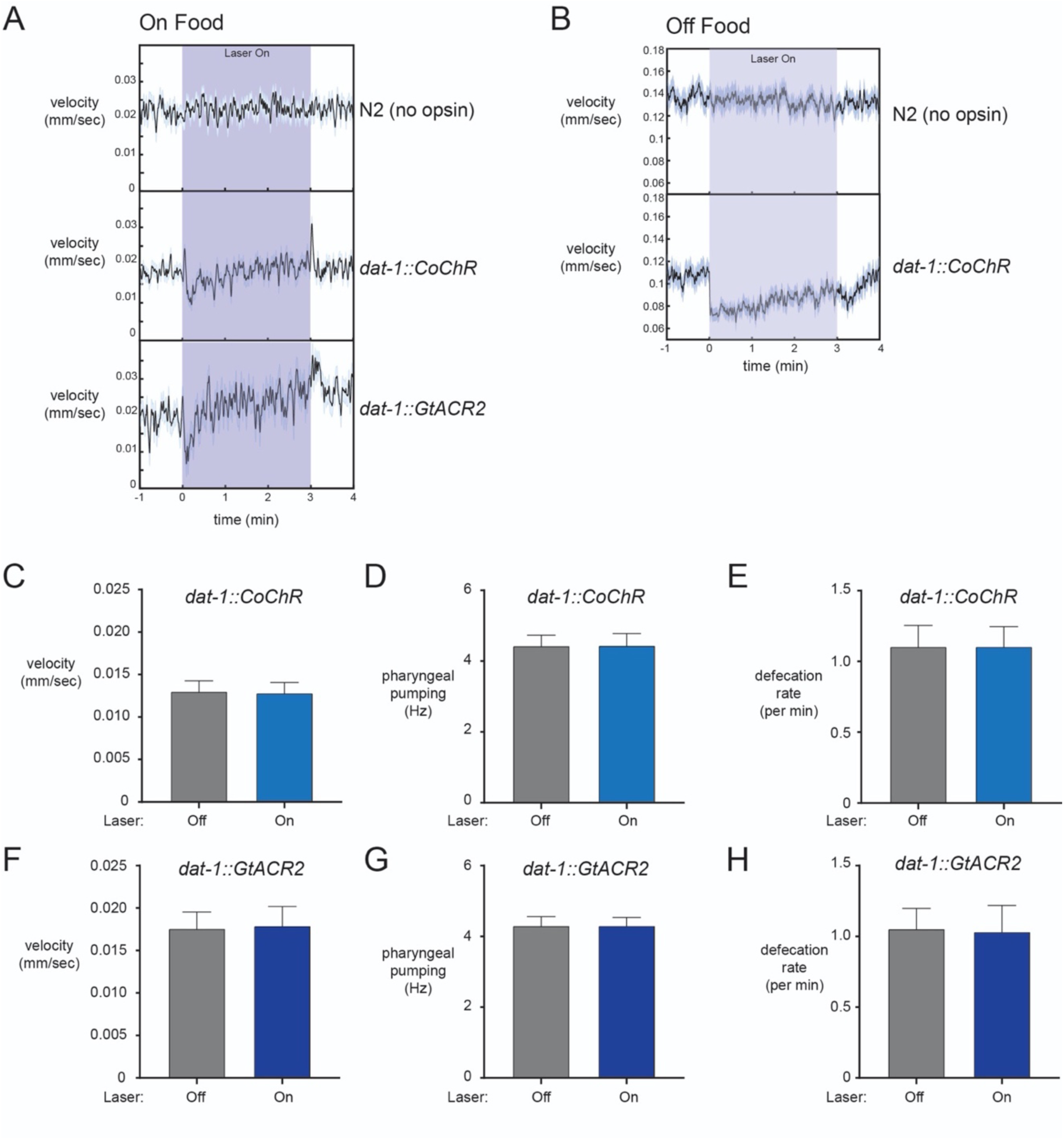
Additional optogenetic studies and analysis. (A) Average velocity during light exposure for wild-type, *dat-1::CoChR*, and *dat-1::GtACR2* animals for experiments conducted in the presence of food. Note that the *dat-1::CoChR* and *dat-1::GtACR2* groups have transient changes in velocity upon light activation and termination. These changes on their own cannot explain egg-laying effects, which are persistent during the lights-on period. n=416-1221 light stimulation events across 12-35 animals for the three genotypes. (B) Effects of *dat-1::CoChR* activation on velocity in the absence of food. Consistent with previous studies, we observe a robust reduction in speed when dopaminergic neurons are activated in the absence of food. WT control shows no effect of the light alone. n=327-385 light stimulation events across 38-44 animals. For (A-B), data are shown as means ± SEM. (C) Average velocity for *dat-1::CoChR* animals during lights-off and lights-on periods. n=35 animals. (D) Average pharyngeal pumping for *dat-1::CoChR* animals during lights-off and lights-on periods. n=35 animals. (E) Average defecation rates for *dat-1::CoChR* animals during lights-off and lights-on periods. n=35 animals. (F) Average velocity for *dat-1::GtACR2* animals during lights-off and lights-on periods. n=12 animals. (G) Average pharyngeal pumping for *dat-1::GtACR2* animals during lights-off and lights-on periods. n=12 animals. (H) Average defecation rates for *dat-1::GtACR2* animals during lights-off and lights-on periods. n=12 animals. For (C-H), data are shown as means ± SEM.

**Figure 5 – Figure Supplement 1.**
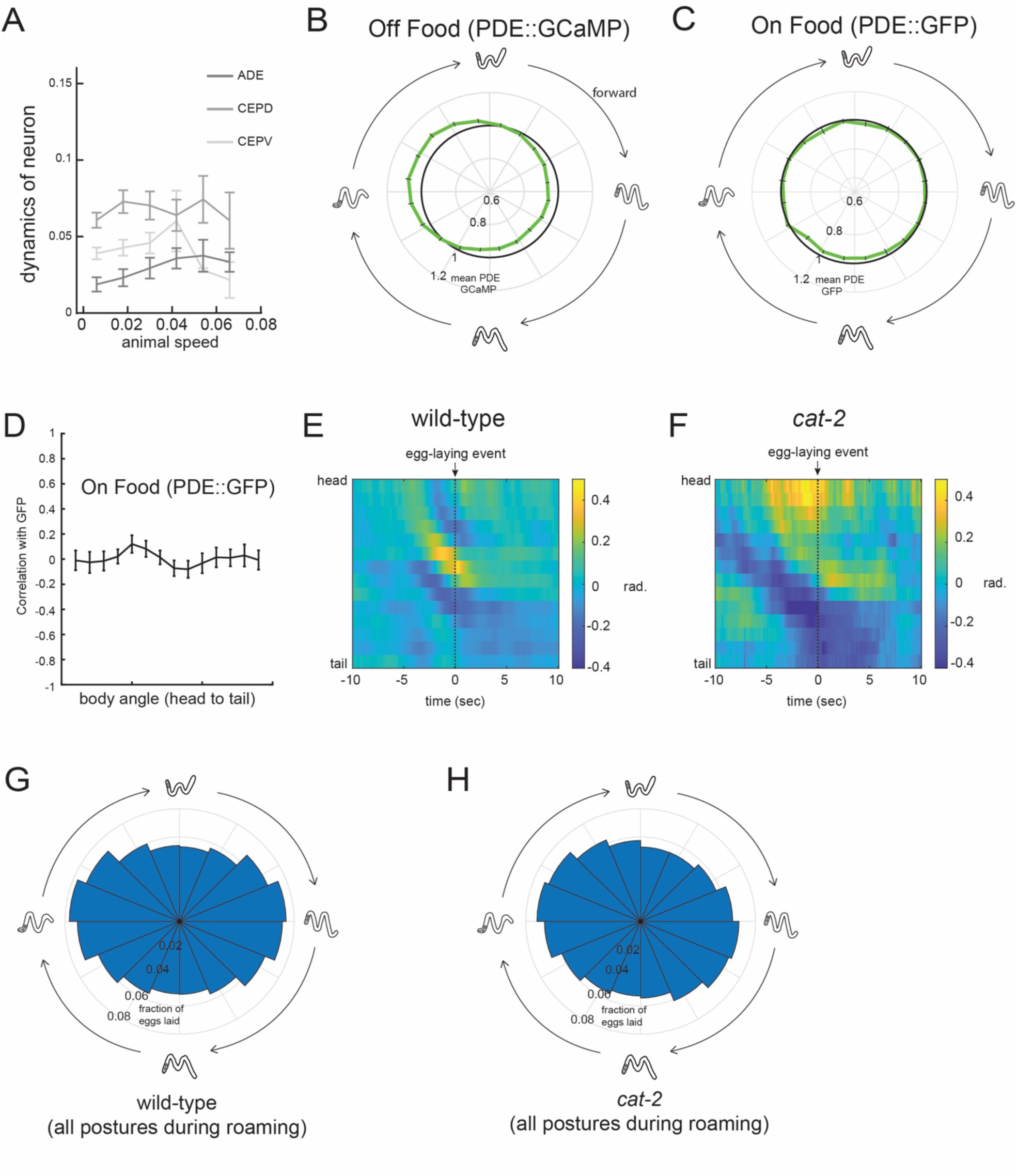
Additional analyses related to PDE activity and egg-laying. (A) Dynamics of ADE, CEPD, and CEPV do not vary with animal speed. Dynamics here is defined as the absolute value of the time derivative of the GCaMP signal for each neuron. n=4-5 animals per condition. (B) Average PDE GCaMP activity during forward-propagating bends for animals recorded in the absence of food. There is significantly reduced activity at the peak phase relative to animals on food (p<0.05, t-test). n=56 animals off food and n=39 animals on food. (C) Average PDE GFP signal during forward-propagating bends for animals recorded in the presence of food. Compare to actual PDE::GCaMP signal in Fig. 5E. n=12. (D) Average correlations of PDE::GFP signal with body curvature. Compare to actual PDE::GCaMP signal in Fig. 5C. n=12. (E) Average posture during roaming egg-laying events depicted as mean of body angles in radians for WT. n = 827 egg-laying events across 30 animals. (F) Average posture during roaming egg-laying events depicted as mean of body angles in radians for *cat-2* mutant animals. n = 140 egg-laying events across 10 animals. (G-H) Histogram depicting phases of the propagating forward bend that wild-type and cat-2 mutant animals display overall, shown with same format as in Fig. 5F-G. Note that there is no significant difference between wild-type and *cat-2*.

